# Cell-intrinsic functions of the transcription factor Bhlhe40 in activated B cells and T follicular helper cells restrain the germinal center reaction and prevent lymphomagenesis

**DOI:** 10.1101/2021.03.12.435122

**Authors:** René Rauschmeier, Annika Reinhardt, Charlotte Gustafsson, Vassilis Glaros, Artem V. Artemov, Reshma Taneja, Igor Adameyko, Robert Månsson, Meinrad Busslinger, Taras Kreslavsky

**Author notes:** these authors contributed equally to the study (René Rauschmeier, Annika Reinhardt). corresponding authors, to whom correspondence should be addressed: Taras Kreslavsky, Meinrad Busslinger.

## Abstract

The generation of high-affinity antibodies against pathogens and vaccines requires the germinal center (GC) reaction – a process that relies on a complex interplay between specialized effector subsets of B and CD4 T lymphocytes – GC B cells and T follicular helper (T_FH_) cells. Intriguingly, several key positive regulators of the GC reaction are common for both cell types. Here, we report that the transcription factor Bhlhe40 is a crucial cell-intrinsic negative regulator affecting both the B and T cell sides of the GC reaction. In activated CD4 T cells, Bhlhe40 was required to restrain proliferation thus limiting the number of T_FH_ cells. In B cells, Bhlhe40 executed its function in the first days after immunization by selectively restricting the generation of the earliest GC B cells but not of early memory B cells or plasmablasts. Conditional Bhlhe40 inactivation confirmed cell-autonomous functions of Bhlhe40 in both GC B and T_FH_ cells, while the GC phenotype was further enhanced upon loss of Bhlhe40 in both cell types. This negative regulation of the GC reaction by Bhlhe40 was of crucial importance, as Bhlhe40-deficient mice with progressing age succumbed to a B cell lymphoma characterized by accumulation of monoclonal GC B-like cells and polyclonal T_FH_ cells in various tissues.

## INTRODUCTION

Generation of high-affinity antibodies in response to pathogens and vaccines depends on a complex interplay between specialized subsets of activated antigen-specific B and T lymphocytes – germinal center (GC) B cells and T follicular helper (T_FH_) cells – that takes place in the course of the GC reaction. Upon activation by cognate antigen, B cells undergo a burst of proliferation and differentiate into plasmablasts (PBs), non-GC-derived memory B cells (MBCs), or GC B cells^1^. The latter cells populate GCs – temporary microanatomical structures that form around follicular dendritic cell (FDC) clusters in B cell follicles of secondary lymphoid organs. The GC reaction is an iterative process that involves repetitive cycles of B cell proliferation, hypermutation of their antibody genes by activation-induced cytidine deaminase (AID), competition of the mutated B cell clones for antigen and subsequent presentation of the processed antigen to a specialized T helper cell subset – T_FH_ cells (reviewed in^2–4^). T cell help provided by T_FH_ cells in response to this antigen presentation is crucial for the survival and expansion of the B cell clones with higher affinity. B cells eventually egress from the GC reaction to give rise to MBCs and high affinity plasma cells (PCs). GCs can be divided histologically into a light zone (LZ) where B cells interact with FDCs that capture and display antigen and T_FH_ cells, and a dark zone (DZ) where proliferation and hypermutation takes place. The GC reaction therefore involves continuous recirculation of B cells between the LZ and the DZ. A wave of expression of the transcription factor Myc that is transiently induced in the LZ proportionally to the magnitude of T cell help^5^ drives proliferation of GC B cells in the DZ^6, 7^, in part through induction of the Myc target gene *Tfap4* encoding the transcription factor AP4^8^. While the GC response is absolutely required for the generation of high-affinity antibodies, active mutagenesis in highly proliferative cells comes at a price, as dysregulation of the GC reaction can lead to lymphomagenesis. Indeed, a wide variety of B cell lymphomas including follicular lymphoma, GC B cell-like diffuse large B cell lymphoma, Burkitt’s lymphoma, Hodgkin’s lymphomas, as well as a number of more rare lymphoma types all originate from the GC reaction_9_.

For reasons that are not well understood, many key regulators of the GC reaction have cell-intrinsic functions in both GC B cells and T_FH_ cells. For example, mice deficient for the transcription factor Bcl6 completely lack GC responses, as Bcl6 is required for the differentiation of both GC B cells and TFH cells^10, 11, 12, 13, 14^. Although the molecular programs regulated by Bcl6 in GC B cells and TFH cells are very different ^15^, in both cell types it functions in part through the direct repression of *Prdm1* that encodes for the transcription factor Blimp1^16^. Blimp1 interferes with the GC B cell molecular program by promoting PC differentiation and shifting the balance of T helper cell differentiation from T_FH_ to TH1 ^1^^7, 18, 19^. The transcription factor Irf4 also promotes PC differentiation when expressed at high levels, however moderate amounts of Irf4 are required for upregulation of Bcl6 during the early phases of the GC response^20, 21^ as well as for T_FH_ differentiation^22^. The GC formation is completely abrogated in absence of the transcription factor Batf^23^ that regulates expression of Bcl6 and Maf^24^ in T_FH_ cells and is required for immunoglobulin class-switch recombination and cell expansion in GC B cells^25 24^. The transcription factor Foxo1 regulates DZ specification in GC B cells^26, 27^ and is required for late-stage GC T_FH_ cell differentiation while inhibiting earlier steps of T_FH_ development^28^. The list of such common regulators is not restricted to transcription factors as mutations in *Rc3h1* encoding the RNA-binding protein Roquin result in a cell-intrinsic expansion of both GC B cells and T_FH_ cells^29, 30^. Thus a number of regulators, many of which are rapidly induced upon lymphocyte activation and therefore are likely to be part of the early response to antigen receptor signaling, play cell-intrinsic roles in both specialized lymphocyte subsets required to mount the GC reaction – GC B cells and T_FH_ cells. Interestingly, while the action of these factors in GC B cells and T_FH_ cells is usually concordant in terms of overall positive or negative effects on the GC response, the exact molecular and cellular functions of these common regulators in the two cell types are often very different.

While a number of positive transcriptional regulators of the GC reaction has been identified to date, little is known about negative regulation of this process. It was previously reported that mice with germline deficiency for the transcription factor Bhlhe40 developed lymphoproliferation and mild autoimmunity with age^31^. The disease was associated with enlarged GCs, but the mechanism behind this phenotype remained uninvestigated. A later report demonstrated that aged *Bhlhe40*^−/−^ mice exhibit decreased regulatory T cell numbers_32_, suggesting one possible explanation for this GC phenotype. In the present study, we aimed to test if Bhlhe40 could function as a cell-intrinsic negative regulator in cells directly participating in the GC reaction. Unexpectedly, this analysis revealed that Bhlhe40 regulates both TFH and GC B cell arms of the GC response in a cell type-specific manner. In B cells, Bhlhe40 restrained the generation of the earliest GC B cells, but not early MBCs or PBs. In activated CD4 T cells, Bhlhe40 restrained proliferation throughout the response, thus limiting T_FH_ cell numbers. With age, a dysregulated GC reaction in *Bhlhe40*^−/−^ mice culminated in the development of a GC B cell lymphoma.

## RESULTS

### A B cell-intrinsic function of Bhlhe40 in restraining the GC reaction

As our RNA-seq experiments (Calderón et al, submitted; also see below) demonstrated that *Bhlhe40* is rapidly upregulated upon B cell activation, we first aimed to test whether Bhlhe40 has a function in activated B cells. Analysis of *Bhlhe40*^−/−^ mice revealed an increased number and size of spontaneous GCs in *Bhlhe40* knockout (KO) mice (Fig. 1a and Supplementary Fig. 1a) and demonstrated the accumulation of cells with GC B and T_FH_ cell phenotypes (Fig. 1b,c and Supplementary Fig. 1b). To test if this phenotype was underlain by a cell-autonomous function of Bhlhe40 in B cells, we established and analyzed WT:*Bhlhe40*^−/−^ mixed bone marrow (BM) chimeras. The overall frequencies of B and T cells originating from each donor closely reflected that observed in BM progenitors (Supplementary Fig. 1c), indicating that Bhlhe40 deficiency does not have a measurable impact on the overall development of T and B lymphocytes. However, the GC B cell compartment in spleens, lymph nodes and Peyer’s patches of these chimeras was dominated by cells originating from the KO donor (Fig. 1d and Supplementary Fig. 1d). This phenotype was independent of the congenic marker used to identify KO cells (Fig. 1d and Supplementary Fig. 1e), indicating that Bhlhe40 deficiency and not allelic variants of genes linked to *Ptprc* (encodes CD45) are responsible for the phenotype. No further increase in GC B cell accumulation was observed when BM from mice double deficient for *Bhlhe40* and its close homolog *Bhlhe41* was used to generate mixed BM chimeras and no GC B cell phenotype was evident when *Bhlhe41*^−/−^ BM was used (Supplementary Fig. 1f), indicating that Bhlhe40, but not Bhlhe41, negatively regulates the GC reaction. As the antigen specificity of ‘spontaneous’ GC B cells observed in mixed BM chimeras is unclear, it was conceivable that their accumulation could have solely reflected some developmental defect (for example in tolerance formation) rather than function of Bhlhe40 in B cell activation. The increase in KO GC B cells in Peyer’s patches (Fig. 1d) suggested that Bhlhe40 may restrain B cell responses to microbiota-derived antigens. To formally test if this transcription factor plays a role in the regulation of responses to foreign antigens, we next immunized mixed BM chimeras with phycoerythrin (PE). PE-specific GC B cells were likewise generated preferentially from KO donor B cells (Fig. 1e). We therefore conclude that Bhlhe40 restrains the GC reaction at least in part through its functions in activated B cells.

**Figure 1.**
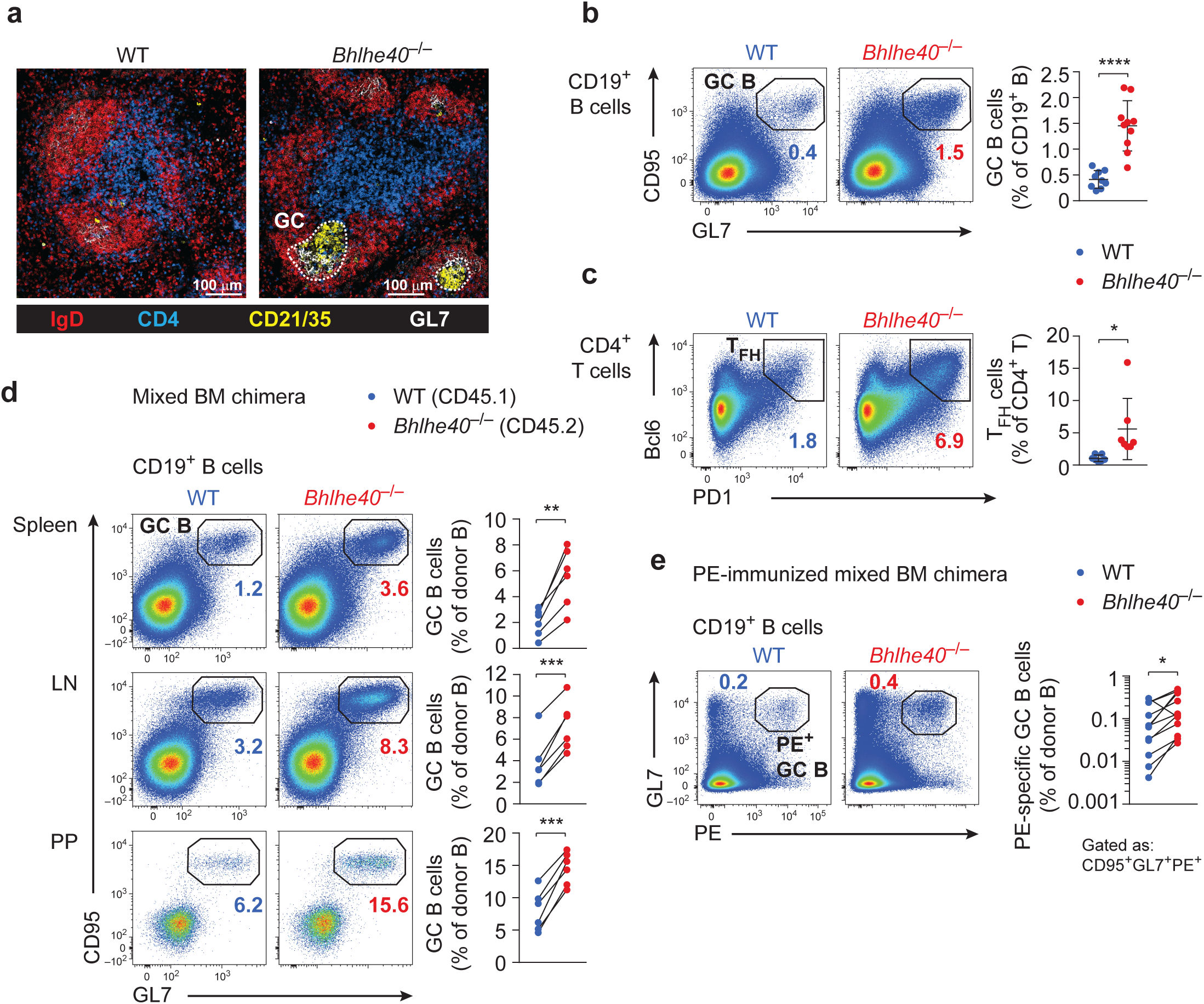
B cell-intrinsic function of Bhlhe40 in restraining the GC reaction. (**a**) Representative images of spleen sections of unchallenged WT and *Bhlhe40*^−/−^ mice as analyzed by confocal immunofluorescence microscopy. Germinal centers (GCs), identified by GL7 (white) and CD21/CD35 (yellow) expression in IgD (red)-negative areas, are highlighted. One of two independent experiments is shown. (**b,c**) Quantification of flow cytometric analysis of GC B cells (**b**) and T_FH_ cells (*c*) in unchallenged WT and *Bhlhe40*^−/−^ mice. Pooled data from two independent experiments with n = 9 (WT) and n = 10 (KO) mice (**b**) and n = 7 mice for each genotype (**c**) are shown. Surface expression of CD95 and GL7 on CD19^+^ B cells (**b**) and surface expression of PD1 and intracellular Bcl6 in CD4^+^ T cells (**c**) are shown. (**d,e**) Mixed BM chimeras were generated by transferring a 1:1 mixture of congenically distinguishable WT (CD45.1^+^) and *Bhlhe40*^−/−^ (CD45.2^+^) BM progenitor cells into lethally irradiated *Rag2*^−/−^ recipients. Mice were analyzed > 6 weeks after transfer. (**d**) Flow cytometric analysis and quantification of the GC B cell population in spleen, peripheral lymph nodes (LN) and Peyer’s Patches (PP) in mixed BM chimeras. One experiment with n = 6 mice is shown, representative of at least three independent experiments. (**e**) Mixed BM chimeras were immunized with PE emulsified in Complete Freund′s adjuvant. Antigen (PE)-specific GC B cells were analyzed by flow cytometry 21 days post immunization. GL7 expression and PE-binding on CD19^+^ B cells (left) and quantification of PE-specific GC B cells (gated as CD95^+^GL7^+^PE^+^) among donor CD19^+^ B cells (right) are shown. Pooled data from three independent experiments with n = 12 mice are shown. Data were analyzed with unpaired Student’s *t*-test (*b*,*c*, mean ± SD is shown) and paired Student’s *t*-test (*d*,*e*). \**P* < 0.05, \*\**P* < 0.01, \*\*\**P* < 0.001, \*\*\*\**P* < 0.0001. One dot represents one mouse.

### Rapid transient upregulation of Bhlhe40 in activated B cells

As a first step to unravel at which timepoint of B cell activation Bhlhe40 could execute its function, we characterized the dynamics of its expression in activated B cells. *In vitro*, *Bhlhe40* was rapidly (maximal level of expression 1 hour after stimulation) but transiently upregulated in B cells upon stimulation with either anti-IgM or anti-CD40 antibodies (with CD40 crosslinking resulting in the highest level of *Bhlhe40* induction), suggesting that both exposure to antigen and interaction with T_FH_ cells could be responsible for *Bhlhe40* induction *in vivo* (Supplementary Fig. 2a). Analysis of data sets from the Immgen database^33^ revealed overall low *Bhlhe40* expression in GC B cells, with a higher level of expression in the light zone (LZ) than in dark zone (DZ) GC B cells (Supplementary Fig. 2b). As several other important regulators of the GC reaction, including Myc and AP4, are also expressed lowly in bulk GC B cells, but are thought to be transiently induced to high levels in a small subset of LZ GC B cells upon interaction with T_FH_ cells and/or antigen^6–8^, we next assessed *Bhlhe40* expression on single-cell level by RNA flow cytometry. Indeed, *Bhlhe40* was expressed by a subset of LZ GC B cells (Supplementary Fig. 2c). Bhlhe40 deficiency, however, did not affect the ratio between LZ and DZ GC B cells (Supplementary Fig. 2d). We conclude that *Bhlhe40* is rapidly but transiently induced upon B cell activation and that its expression in a subset of LZ GC B cells may likewise reflect transient induction upon interaction with antigen and/or T_FH_ cells.

### Bhlhe40 restricts the generation of the earliest GC B cells

The expression analysis described above suggested that Bhlhe40 could execute its function early upon B cell activation after the initial interaction with antigen and/or T_FH_ cells as well as later in the response, during the course of the ongoing GC reaction, through its transient induction in the LZ. To start addressing these possibilities, we aimed to test at which point of the response Bhlhe40-deficient B cells gain their competitive advantage. To have sufficient cell numbers for characterization of the earliest stages of the response, we crossed *Bhlhe40*^−/−^ mice to B1-8^hi^ *Igh* knock-in mice in which B cells with λ light chains (Igλ) recognize the hapten 4-hydroxy-3-nitrophenylacetyl (NP)^34^. We then co-transferred congenically distinguishable *Bhlhe40*^−/−^*Igh*^B1-8hi/+^ and *Bhlhe40*^+/+^*Igh*^B1-8hi/+^ splenocytes (with confirmed 1:1 ratio of Igλ^+^ B cells) into WT recipients followed by NP-OVA immunization (Fig. 2a). In line with observations made for the polyclonal responses to PE (Fig. 1e), these experiments demonstrated that the majority of donor-derived GC B cells were originating from the KO donor (Fig. 2b,c). The competitive advantage of Bhlhe40- deficient cells was evident for the first GC B cells that emerged around day 3 after immunization, reached its maximum by day 6 after immunization and remained stable thereafter (Fig. 2c). These results suggested that Bhlhe40 restrains the earliest stages of the GC reaction and Bhlhe40 expression in B cells may become dispensable in mature GCs. We therefore focused our attention on the events that take place during the emergence of the first GC B cells around day 3-4 after immunization. At this point, activated Igλ^+^ B1-8^hi^ B cells formed three distinct populations: CD138^+^ early PBs, a CCR6^+^GL7^lo/int^ population likely including non-GC-derived early MBCs as well as common precursors for all activated populations^1, 35, 36^ and CCR6^−^GL7^hi^ early GC B cells (Fig. 2b). Strikingly, while in the competitive transfer experiments KO B1-8^hi^ B cells exhibited a clear advantage in the GC B cell compartment, ratios in both PBs and in the CCR6^+^ activated B cell population closely reflected the ratio in Igλ^+^ B cells at the time of the transfer (Fig. 2b). Of note, no difference in Bcl6 expression level was observed for WT and KO GC B cells (Supplementary Fig. 3a). The phenotype was the same when congenic markers on WT and KO B1-8^hi^ B cells were inverted (Supplementary Fig. 3b). Taken together, these results demonstrated that Bhlhe40 is a highly selective negative regulator of the earliest GC B cells but not of other subsets of activated B cells.

**Figure 2.**
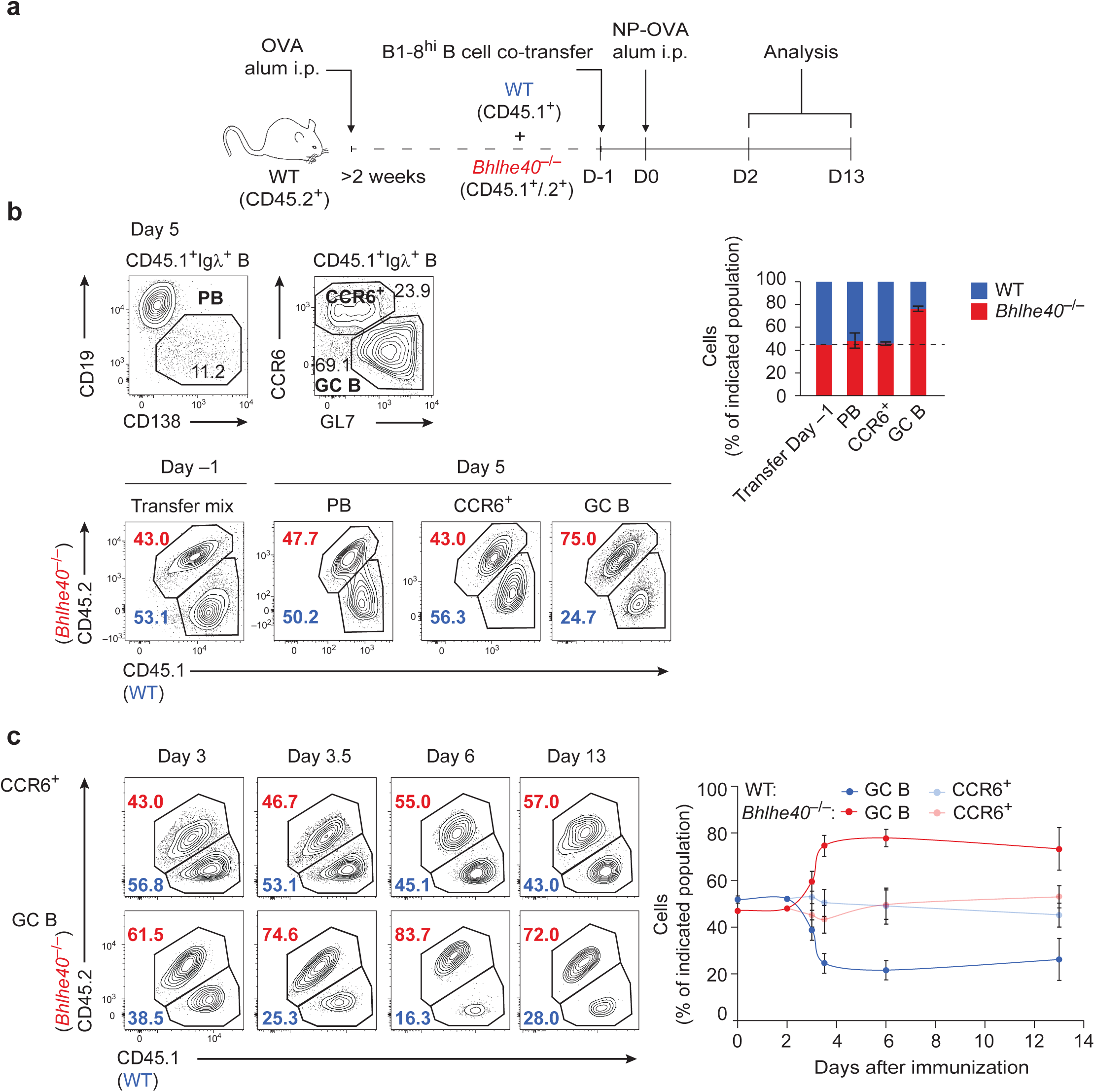
Bhlhe40 restricts the earliest stages of the GC B cell response. (**a**) Schematic representation of the experimental setup for **b-c**. OVA-primed CD45.2^+^ WT mice were injected with a 1:1 mixture of congenically distinguishable WT (CD45.1^+^) and *Bhlhe40*^−/−^ (CD45.1^+^/CD45.2^+^) B1-8^hi/+^ splenocytes and were immunized with NP-OVA in alum next day. (**b**) (top) Surface expression of CD138, CCR6 and GL7 on CD45.1^+^Igλ^+^CD19^+^ B cells (intracellular Igλ staining for PB gating) and quantification of WT and *Bhlhe40*^−/−^ -derived cells among total CD45.1^+^Igλ^+^CD19^+^ B cells at the time of the transfer, as well as among plasmablasts (PB), CCR6^+^ and GC B cells on day 5 post immunization is shown. One experiment with n = 4 mice is shown, representative of three independent experiments. (*c*) Representative flow cytometric analysis and quantification of WT and *Bhlhe40*^−/−^-derived cells among CD45.1^+^Igλ^+^CD19^+^ CCR6^+^ and GC B cells at indicated time points post immunization. Gating for total Igλ^+^CD19^+^ B cells was applied for day 0 (transfer mix) and day 2 (prior to GC B cell emergence) after immunization. The time course is based on pooled data from two independent experiments with n = 4-16 mice for each time point. One dot represents the mean with SD for each time point.

As Bhlhe40 is a negative regulator of proliferation in several cell types^37, 38^, we next tested if Bhlhe40 deficiency affects proliferation of GC B cells or the CCR6^+^ activated B cell population, focusing these experiments on the timepoint when the very first GC B cells were detectable in most of the mice. Incorporation of EdU by B1-8^hi^ WT and KO B cells was comparable, and GC B cells of both genotypes exhibited much higher frequencies of EdU^+^ cells than the CCR6^+^ population (Fig. 3a). Expectedly, no difference in EdU incorporation was detectable at later timepoints (data not shown) or in ‘spontaneous’ GC B cells in mixed BM chimeras (Supplementary Fig. 4a). Proliferation of *in vitro* activated B cells was also comparable between co-cultured WT and KO B cells across a range of anti-IgM and anti-CD40 concentrations, and the ratio between WT and KO B cells remained stable in these cultures (Supplementary Fig. 4b), suggesting that, at least *in vitro*, survival of B cells is also not affected by Bhlhe40 deficiency. In line with this notion, Bhlhe40 deficiency did not result in any measurable change in the frequency of cleaved Caspase 3-positive B1-8^hi^ B cells *in vivo*, and very little apoptosis took place in GC B cells at these early timepoints irrespectively of their genotype (Fig. 3b). Likewise, apoptosis of polyclonal GC B cells was not affected by Bhlhe40 deficiency in mature GCs in mixed BM chimeras (Supplementary Fig. 4c).

**Figure 3.**
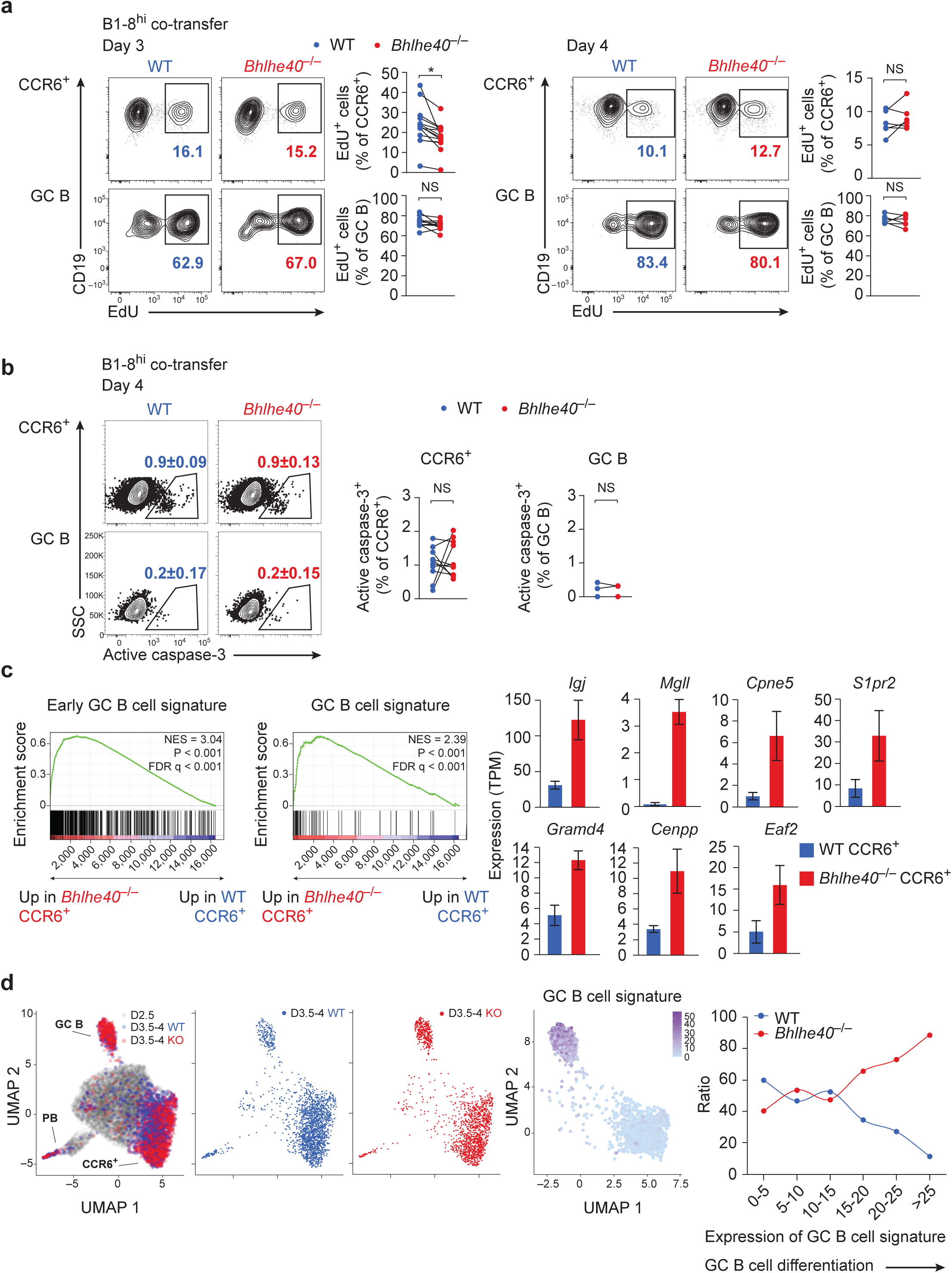
Bhlhe40 restrains the generation of early GC B cells. (**a-d**) OVA-primed CD45.2^+^ WT mice were injected with a 1:1 mixture of congenically distinguishable WT (CD45.1^+^) and *Bhlhe40*^−/−^ (CD45.1^+^/CD45.2^+^) (**a** (day 3),*c*,*d*) or WT (CD45.1^+^/CD45.2^+^) and *Bhlhe40*^−/−^ (CD45.1^+^) (**a** (day 4),*b*) B1-8^hi/+^ splenocytes and were immunized with NP-OVA in alum next day. (*a*,*b*) Flow cytometric analysis and quantification assessing the proliferation (*a*) and frequency of apoptotic cells (*b*) of WT and *Bhlhe40*^−/−^ CCR6^+^ and GC B cells on day 3 (**a**) and 4 (**a,b**) post immunization. (**a**) EdU was i.v. injected 4 and 2 hours prior to analysis. (*b*) Apoptotic cells were identified by intracellular staining for active caspase-3. (**a**) One experiment with n = 14 (day 3) and n = 6 (day 4) mice is shown. Representative of at least two independent experiments. (**b**) Pooled data from two independent experiments with n = 10 mice are shown. Note that only in 2 mice out of 10 active caspase-3^+^ cells were detectable in the GC B cell compartment. Flow cytometry plots show electronically merged data from 3 mice, and mean ± SD is shown. Data were analyzed with paired Student’s *t*-test. \**P* < 0.05. One dot represents one mouse. (**c**) Left – Gene set enrichment analysis (GSEA) on an RNA-seq dataset ranked by KO/WT fold change in day 4 CCR6^+^ B1-8^hi^ B cells using a published GC B cell signature^39^ (‘GC B cell signature’) or the top 500 genes upregulated in WT day 4 B1-8^hi^ GC B cells when compared to WT day 4 CCR6^+^ B cells (‘Early GC B cell signature’) as gene sets. NES, normalized enrichment score; FDR, false-discovery rate. Right – examples of genes that were enriched in KO CCR6^+^ B1-8^hi^ B cells as identified by GSEA. Expression in WT and *Bhlhe40*^−/−^ CCR6^+^ B cells as assessed by RNA-seq analysis on day 3.5-4 post immunization is shown. (**d**) scRNA-seq comparison of WT and KO GC B cell differentiation. Left – UMAP plots showing the distribution of WT (days 2.5 and/or 3.5-4 after immunization) and KO (days 3.5-4 after immunization) B1-8^hi^ B cells. Middle – UMAP plot showing the expression of GC B cell signature genes in clusters that were included for the subsequent WT/KO comparison: the frequency of WT (blue) and KO (red) B cells among cells binned by the strength of expression of GC B signature genes is shown.

The results described above suggested that Bhlhe40 is unlikely to regulate the size of the GC B cell compartment through effects on apoptosis or proliferation. We therefore hypothesized that Bhlhe40 could act through limiting the differentiation of activated cells to the GC B cell ‘lineage’. To find evidence for or against this hypothesis, we next performed RNA-seq on double-sorted WT and KO CCR6^+^ and GC B cell populations from the B1-8^hi^ co-transfer system described above at day 4 after immunization. This analysis revealed very limited changes in gene expression between WT and KO cells for both populations with less than a dozen genes significantly (adjusted *P* value < 0.05, fold change ≥ 2) changing their expression in the KO cells (Supplementary Fig. 5a). None of these genes could be obviously connected to the phenotype. Unexpectedly, while CCR6^+^ cells expressed *Bhlhe40*, virtually no expression was observed in these early GC B cells (Supplementary Fig. 5b), suggesting that *Bhlhe40* functions in precursors of early GC B cells likely within the CCR6^+^ activated B cell population. Indeed, gene set enrichment analysis (GSEA) demonstrated that a GC B cell signature gene set^39^ was mildly but significantly upregulated in KO CCR6^+^ B cells compared to their WT counterparts (Fig. 3c). Likewise, GSEA using top 500 genes that were expressed higher in day 4 WT GC B cells than in WT CCR6^+^ activated B cells (‘Early GC B cell signature’) revealed upregulation of this gene set in the KO CCR6^+^ population (Fig. 3c). To test if Bhlhe40 may directly repress some of these GC B cell signature genes, we performed ChIP-seq detection of the genome-wide Bhlhe40 binding in naïve B cells stimulated *in vitro* for 4 hours with anti-CD40 antibody. This analysis demonstrated enrichment of the expected CTCGTG E-box motif in Bhlhe40 peaks, revealed the preferential promoter and gene body binding of Bhlhe40 (Supplementary Fig. 5c), and suggested that some of the GC B cell signature genes upregulated in the KO cells may indeed be directly repressed by Bhlhe40 (Supplementary Fig. 5d).

We next compared WT and KO Igλ^+^ B1-8^hi^ B cells from days 3.5 and 4 (timepoints pooled) after immunization by single cell (sc)RNA-seq. An additional WT dataset from day 2.5 after immunization was included in the initial analysis to better define the population structure of early B cell activation and capture developmental transitions. Comparison of WT and KO cells was then performed using cells from day 3.5-4 only. As described in detail elsewhere (Glaros et al, submitted), this analysis confirmed that activated B cells at day 3.5-4 formed three distinct groups corresponding to GC B cells, PBs and the CCR6^+^ population containing activated precursors and early MBCs (Supplementary Fig. 5e). Both WT and KO cells were present across all the populations, but as expected, KO cells were enriched among GC B cells (Fig. 3d). Similar to the bulk RNA-seq results, scRNA-seq revealed only minor changes in gene expression between WT and KO populations (data not shown). To test if Bhlhe40 influences GC B cell differentiation, we next ranked cells by their level of expression of GC B cell signature genes (Fig. 3d, Supplementary Fig. 5f) and assessed the ratio between WT and KO cells along this trajectory. This analysis revealed a gradual increase in the relative frequency of KO cells in the course of this early GC B cell development (Fig. 3d), indicating that Bhlhe40-deficient B cells gained a competitive advantage at the earliest steps of GC B cell differentiation and suggesting their possible accelerated progression along this differentiation trajectory. Taken together, these results suggest that Bhlhe40 restrains the generation of the earliest GC B cells, that Bhlhe40-deficient cells gain their competitive advantage gradually in the course of early GC B cell differentiation, and that Bhlhe40 may function through mild repression of some GC B cell program genes in early activated B cells.

### A T cell-intrinsic function of Bhlhe40 restrains the T_FH_ cell response

The results described above clearly identified a cell-intrinsic function of Bhlhe40 in activated B cells. To test if increased GC activity in *Bhlhe40*^−/−^ mice can be fully explained by its role in B lymphocytes, we sought to inactivate *Bhlhe40* selectively in this cell type. To this end, we generated a new conditional allele of *Bhlhe40* (Supplementary Fig. 6a) and crossed mice carrying this allele to a panel of Cre lines. Ablation of *Bhlhe40* in all hematopoietic cells with *Vav1*-Cre phenocopied steady-state GC B and T_FH_ cell phenotypes of *Bhlhe40*^−/−^ mice, indicating that they can be fully explained by Bhlhe40 function in the hematopoietic system (Fig. 4a). Unexpectedly, while deletion of *Bhlhe40* in B cells with *Mb1*-Cre (active from pro-B cell stage on) also resulted in a GC B and T_FH_ cell increase, both phenotypes were markedly less pronounced than in *Vav1*-Cre *Bhlhe40*^fl/fl^ and *Bhlhe40*^−/−^ mice (Fig. 4b), despite the equally high deletion efficiencies for both Cre lines in B cells (Supplementary Fig. 6b). Inactivation of *Bhlhe40* later in B cell development, in immature B cells, with *Cd23*-Cre replicated the phenotype observed with *Mb1*-Cre deletion (Fig. 4c), while no phenotype was observed in *Aicda*-Cre *Bhlhe40*^fl/fl^ mice (Fig. 4d). As *Aicda*-Cre is predominantly active in GC B cells that express the highest level of *Aicda* (encodes AID), these results are in line with the notion that Bhlhe40 functions in a narrow time window very early in GC B cell differentiation and that it is dispensable in mature GC B cells. The discrepancy in the magnitude of the phenotypes observed in B cell-specific (Fig. 4b,c) and pan-hematopoietic (Fig. 4a) *Bhlhe40* knockouts indicated that inactivation of Bhlhe40 in another cell type in the hematopoietic system may contribute to the overall phenotype in *Bhlhe40*^−/−^ and *Vav1*-Cre *Bhlhe40*^fl/fl^ mice. In search for this additional cell type, we next deleted *Bhlhe40* in T cells with *Cd4*-Cre. Strikingly, this also resulted in a measurable increase in both T_FH_ cell and GC B cell compartments (Fig. 4e). In line with the notion that T cell help availability is a limiting factor in the GC reaction^40^, both phenotypes were stronger than those observed in case of B cell-specific deletion, with the T_FH_ phenotype approximating that of *Vav1*-Cre *Bhlhe40*^fl/fl^ mice. Thus, Bhlhe40 restrains the GC reaction through its cell-intrinsic functions both in B and T lymphocytes.

**Figure 4.**
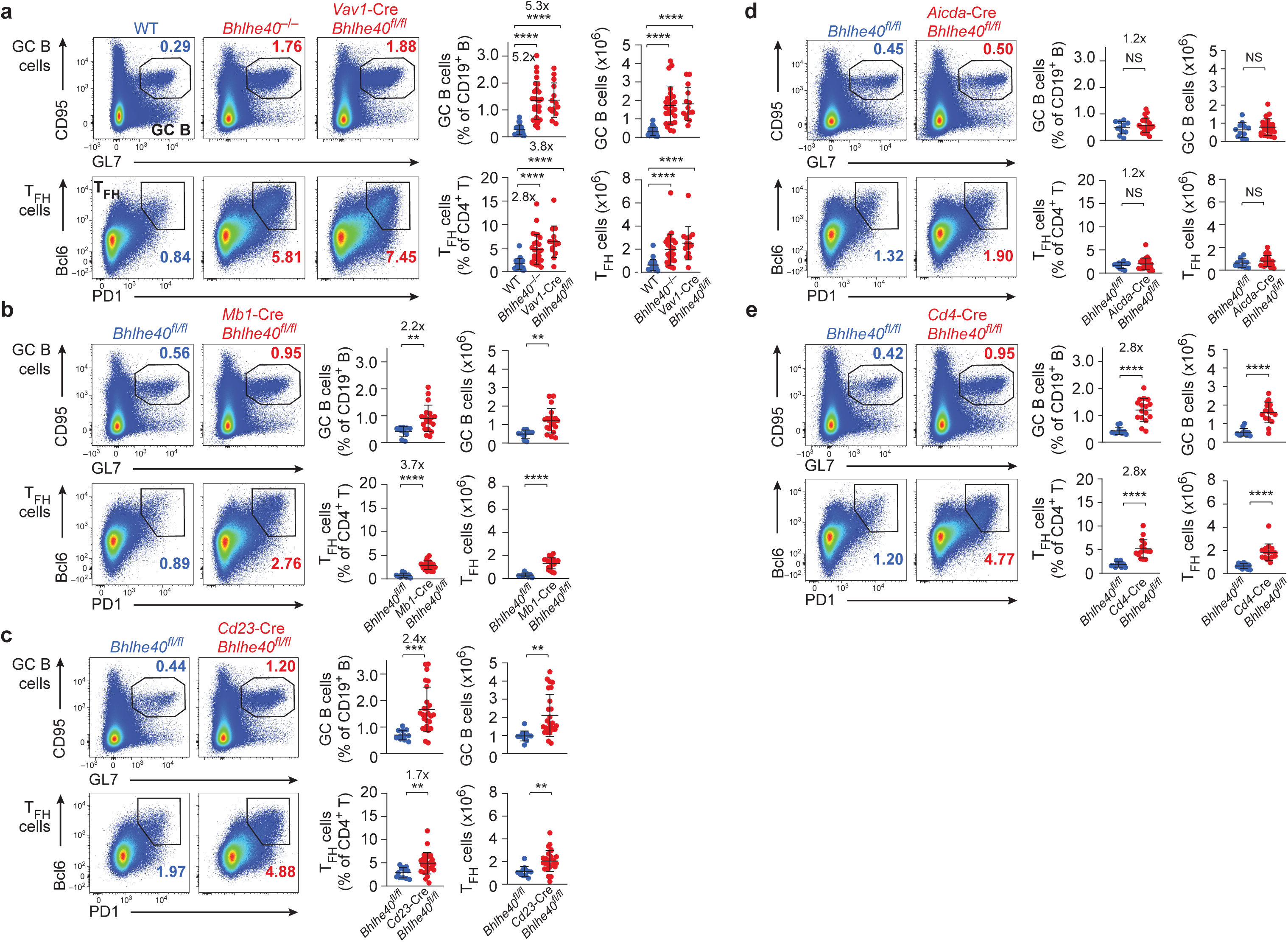
Bhlhe40 restrains the GC reaction through its cell-intrinsic functions in both B and T cells. Flow cytometric analysis and quantification of GC B cells and T_FH_ cells in unchallenged (**a**) WT, *Bhlhe40*^−/−^ and *Vav1*-Cre *Bhlhe40*^fl/fl^ mice, (**b**) *Bhlhe40*^fl/fl^ and *Mb1*-Cre *Bhlhe40*^fl/fl^ mice, (**c**) *Bhlhe40*^fl/fl^ and *Cd23*-Cre *Bhlhe40*^fl/fl^ mice, (**d**) *Bhlhe40*^fl/fl^ and *Aicda*-Cre *Bhlhe40*^fl/fl^ mice, (**e**) *Bhlhe40*^fl/fl^ and *Cd4*-Cre *Bhlhe40*^fl/fl^ mice. Pooled data from three independent experiments with (**a**) n = 24 (WT), n = 27 (*Bhlhe40*^−/−^), n = 14 (*Vav1*- Cre *Bhlhe40*^fl/fl^), (**b**) n = 10 (*Bhlhe40*^fl/f^) and n = 19 (*Mb1*-Cre *Bhlhe40*^fl/fl^), (**c**) n = 12 (*Bhlhe40*^fl/f^) and n = 26 (*Cd23*-Cre *Bhlhe40*^fl/fl^), (**d**) n = 11 (*Bhlhe40*^fl/f^) and n = 22 (*Aicda*-Cre *Bhlhe40*^fl/fl^), and (**e**) n = 11 (*Bhlhe40*^fl/f^) and n = 16 (*Cd4*-Cre *Bhlhe40*^fl/fl^) mice are shown. Fold changes are indicated. Mean values are shown with SD and data were analyzed with Student’s *t*-test. One dot represents one mouse. \**P* < 0.05, ** *P* < 0.01, *** *P* < 0.001, **** *P* < 0.0001.

We next aimed to test if Bhlhe40 executes its function upon T cell activation. In line with this possibility, *Bhlhe40* was previously shown to be rapidly induced upon TCR crosslinking in naïve T cells^32, 41^ and was expressed in T_FH_ cells (Fig. 5a). To further characterize the function of Bhlhe40 in CD4 T cells, we once again took advantage of the WT:*Bhlhe40*^−/−^mixed BM chimera approach. As for GC B cells, ‘spontaneous’ T_FH_ cells in mixed BM chimeras were preferentially generated from KO cells (Fig. 5b, Supplementary Fig. 7a). A less pronounced increase was also observed in non-T_FH_ activated CD4 T cells but the frequency of T_FH_ cells from activated (CD44^hi^CD62L^lo^) cells remained higher for KO CD4 T cells (Fig. 5b). These results suggest that Bhlhe40 expression has a negative impact on the activated CD4 T cell compartment, and this impact is stronger in the case of the T_FH_ cell subset. To test if this phenotype can be recapitulated in response to a foreign antigen, we next immunized mixed BM chimeras with the peptide antigens Ag85b_280-294_ or 2W1S in complete Freund’s adjuvant (CFA). Quantification of the responding cells from both donors with MHC class II tetramers revealed a significant increase in the frequency of antigen-specific CD4 T cells derived from the KO donor cells (Fig. 5c and Supplementary Fig. 7b) and the vast majority of these cells had a PD1^+^CXCR5^+^ T_FH_ cell phenotype (Fig. 5c). We conclude that in addition to its function in B cells, Bhlhe40 restrains the T_FH_ cell response in a cell-intrinsic manner.

**Figure 5.**
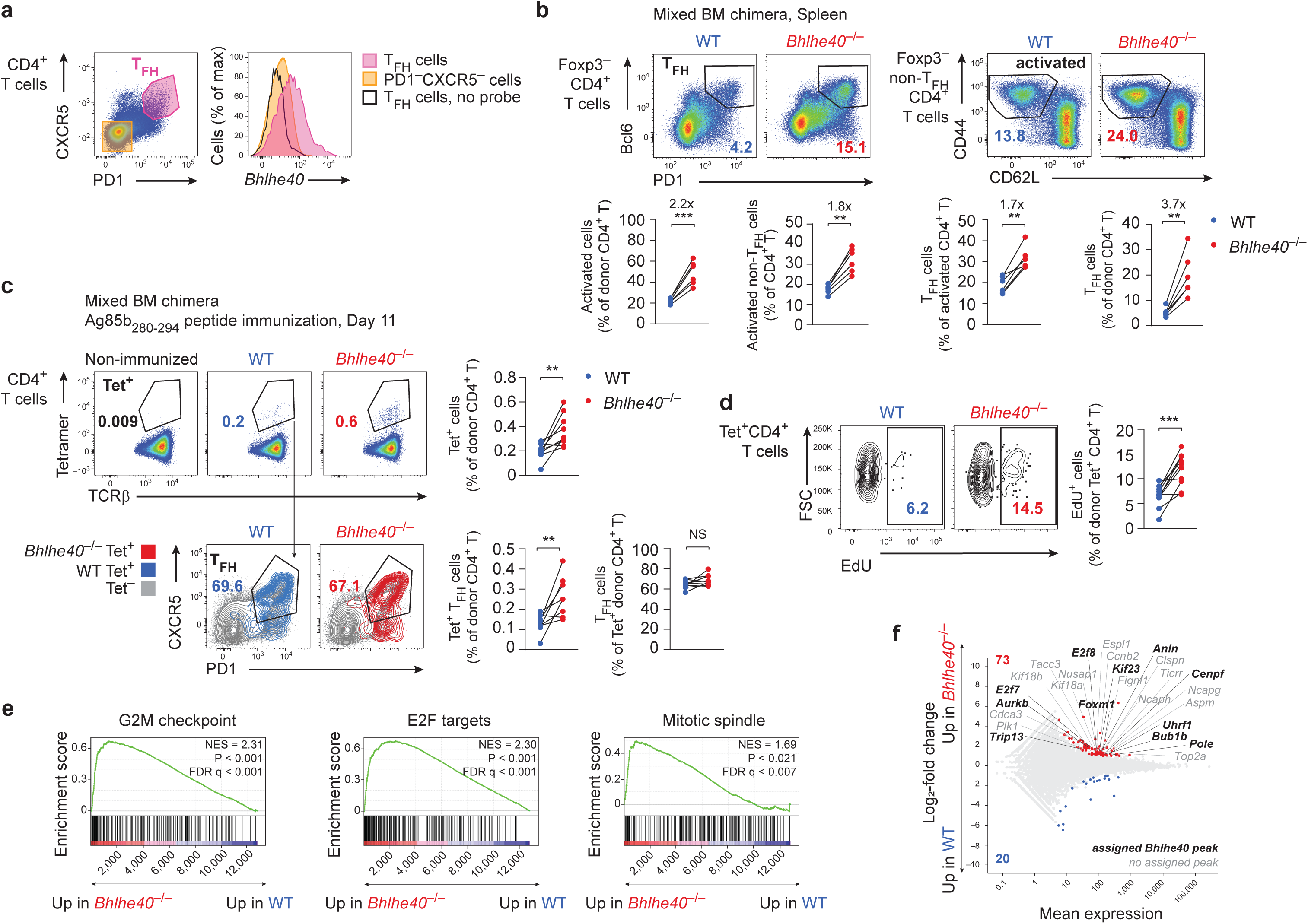
Bhlhe40 limits the proliferation of T_FH_ cells. (**a**) *Bhlhe40* expression in TFH cells among CD4^+^ T cells from spleens of unchallenged WT mice as analyzed by RNA flow cytometry. One experiment with pooled data from n = 2 mice is shown. Representative of two independent experiments. (**b**) Mixed BM chimeras were generated by transferring a 1:1 mixture of congenically distinguishable WT (CD45.1^+^) and *Bhlhe40*^−/−^ (CD45.2^+^) BM progenitor cells into lethally irradiated *Rag2*^−/−^ recipients. Mice were analyzed > 6 weeks after transfer. Expression of PD1 and Bcl6 by WT and *Bhlhe40*^−/−^ Foxp3^−^CD4^+^ T cells (top row left), flow cytometric analysis of activated CD44^+^CD62L^−^ cells among WT and *Bhlhe40*^−/−^ Foxp3^−^non-T_FH_ CD4^+^ T cells (top row right), and quantification among total or activated WT and *Bhlhe40*^−/−^ Foxp3^−^CD4^+^ T cells in the spleen (bottom row) are shown. For quantification, T_FH_ cells were gated as Foxp3^−^Bcl6^+^CD4^+^ T cells or Foxp3^−^Bcl6^+^PD1^+^CD4^+^ T cells (in ′T_FH_ of donor CD4^+^ T′). One experiment with n = 6 mice is shown. Representative of two independent experiments. (*c*-*f*) Mixed BM chimeras were generated by transferring a 1:1 mixture of congenically distinguishable WT (CD45.1^+^/CD45.2^+^) and *Bhlhe40*^−/−^ (CD45.1^+^) BM progenitor cells into lethally irradiated WT (CD45.2^+^) recipients. Mice were immunized with *M. tuberculosis* Ag85b_280-294_ peptide emulsified in Complete Freund′s adjuvant > 7 weeks after transfer. (**c**) Flow cytometric analysis and quantification of Ag85b-tetramer-binding CD4^+^ T cells and T_FH_ cells among WT and *Bhlhe40*^−/−^ CD4^+^ T cells in the spleen on day 11 post immunization. (**d**) Immunized mixed BM chimeras were i.p. injected with EdU 18 and 14 hours before harvest on day 11. EdU incorporation by Ag85b-tetramer-binding WT and *Bhlhe40*^−/−^ CD4^+^ T cells is shown. One experiment with n = 9 (*c*) and n = 10 (**d**) mice is shown, respectively. Representative of two independent experiments each. Data were analyzed with paired Student’s *t*-test. \**P* < 0.05, \*\**P* < 0.01, \*\*\**P* < 0.001. One dot represents one mouse. (**e,f**) RNA-seq analysis of Ag85b-tetramer-binding WT and *Bhlhe40*^−/−^ CD4^+^ T cells isolated from mixed BM chimeras 11 days post immunization. (**e**) GSEA showing top 3 most enriched gene sets from MSigDB Hallmark collection: G2M checkpoint, E2F target and mitotic spindle genes depicting the change in their expression in *Bhlhe40*^−/−^ relative to that in WT Ag85b-tetramer-binding CD4^+^ T cells. NES, normalized enrichment score; FDR, false-discovery rate. (**f)** Comparison of changes in gene expression induced by Bhlhe40 deficiency in Ag85b-tetramer-binding CD4^+^ T cells. Log2-transformed *Bhlhe40*^−/−^/WT fold changes are plotted and the number of significantly (> 2-fold, adjusted *P* < 0.05) up- and downregulated genes is indicated. Cell cycle-related genes (GO:0007049) that significantly (> 2-fold, adjusted *P* < 0.05) changed their expression in *Bhlhe40*^−/−^ cells are highligted. Gene names in bold correspond to genes with assigned Bhlhe40 ChIP-seq peaks

### Bhlhe40 restrains TFH cell proliferation

As Bhlhe40 and its close homolog Bhlhe41 were reported to have moderate anti-proliferative effects in a number of cell types^37^, we next tested if the competitive advantage of Bhlhe40-deficient antigen-specific CD4 T cells in mixed BM chimeras was underlain by their increased proliferation. Indeed, tetramer-positive KO CD4 T cells showed significantly increased EdU incorporation when compared to their WT counterparts (Fig. 5d). RNA-seq comparison of WT and KO tetramer-positive cells revealed upregulation of signatures associated with cell cycle (G2M checkpoint, E2F target genes, and mitotic spindle) in the KO T cells (Fig. 5e, f). At least some of these genes (*E2f7*, *E2f8, Foxm1*, *Aurkb* and many others) are likely to be directly repressed by Bhlhe40 in WT T cells as evidenced by the binding of Bhlhe40 in the proximity of these genes (Fig. 5f and Supplementary Fig. 7c) in a ChIP-seq dataset from *in vitro* activated CD4 T cells^42^. More genes were upregulated than downregulated in the KO cells (73 vs 20 genes with adjusted *P* value < 0.05, fold change ≥ 2), in line with the notion that Bhlhe40 functions predominantly as a transcriptional repressor^37^. Importantly, no change in the expression of known positive or negative regulators of T_FH_ cell development and function (such as *Bcl6*, *Prdm1*, *Irf4*, *Maf*, *Batf*, *Foxo1,* or *Rc3h1*) was observed (Supplementary Fig. 7d). As Bhlhe40 recently emerged as an important regulator of cytokine production by different T cell subsets^42–45^, we also checked the expression of genes encoding these cytokines in our T_FH_-dominated antigen-specific T cells, but did not observe any changes in their expression in the KO cells (Supplementary Fig. 7d). We conclude that Bhlhe40 negatively regulates T_FH_ cell expansion, at least in part through the direct repression of cell cycle-related genes.

### Lymphomagenesis in Bhlhe40-deficent mice

GCs are thought to be the site of origin of the majority of mature human B cell lymphomas^9^. We therefore next tested if dysregulation of multiple aspects of the GC reaction in *Bhlhe40*^−/−^ mice could ultimately lead to lymphomagenesis. Indeed, by 20 months of age nearly all tested *Bhlhe40*^−/−^ mice succumbed to a fatal disease manifested by the formation of ectopic lymphocyte-containing nodules in a variety of organs (Fig. 6a-c) as well as by splenomegaly. While splenomegaly was also observed in previous studies and was attributed to autoimmunity-associated lymphoproliferation^31, 32^, ectopic lymphocyte nodules were not reported to date, likely because younger mice were analyzed in these earlier studies. Flow cytometry revealed a heterogeneous cellular composition of these nodular structures with the presence of CD4 and CD8 T cells as well as B cells (Fig. 6d). Strikingly, the majority of these B cells had a GC B cell phenotype, and a large fraction of CD4 T cells expressed markers of T_FH_ cells (Fig. 6e). In line with their likely GC origin, both populations expressed Bcl6 (Fig. 6e), and GC B-like cells expressed *Aicda* (Supplementary Fig. 8a), however the GC architecture was not preserved in the nodules and FDC clusters were absent (Fig. 6f). T_FH_- like cells in these structures were highly polyclonal (Fig. 6g), while GC B-like cells, in stark contrast to their normal WT or *Bhlhe40*-deficient counterparts, were dominated by single clones as evidenced by the predominant utilization of single IgH and IgΚ V segments revealed by RNA-seq (Fig. 6h). Interestingly, one of the two tumors subjected to RNA-seq expressed the *Igha* constant region while the second lymphoma expressed *Ighm* (Supplementary Fig. 8b). These results are consistent with a model in which dysregulation of the GC reaction in *Bhlhe40*^−/−^ mice with age results in malignant transformation of a B cell clone, while an expanded T_FH_ cell compartment may represent a component of the tumor ‘stroma’. Of note, a similar cellular composition in tumors, with transformed B cells being outnumbered by T cells, has been reported for several human GC-derived B cell lymphomas^46, 47^. To test if these B cells are indeed malignantly transformed and if they depend on the presence of T_FH_-like cells, we transferred sorted B and CD4 T cells from diseased mice separately or in combination into *Rag2*^−/−^ recipients. These experiments demonstrated that B cells were required and sufficient to transfer the disease (Fig. 6i) and the presence of T_FH_-like cells did not accelerate its progression, indicating that, at least at this late stage, the transformed GC B-like cells do not depend on T cell help. We conclude that the dysregulation of the GC reaction observed in Bhlhe40-deficient mice culminates in the formation of a GC-derived B cell lymphoma.

**Figure 6.**
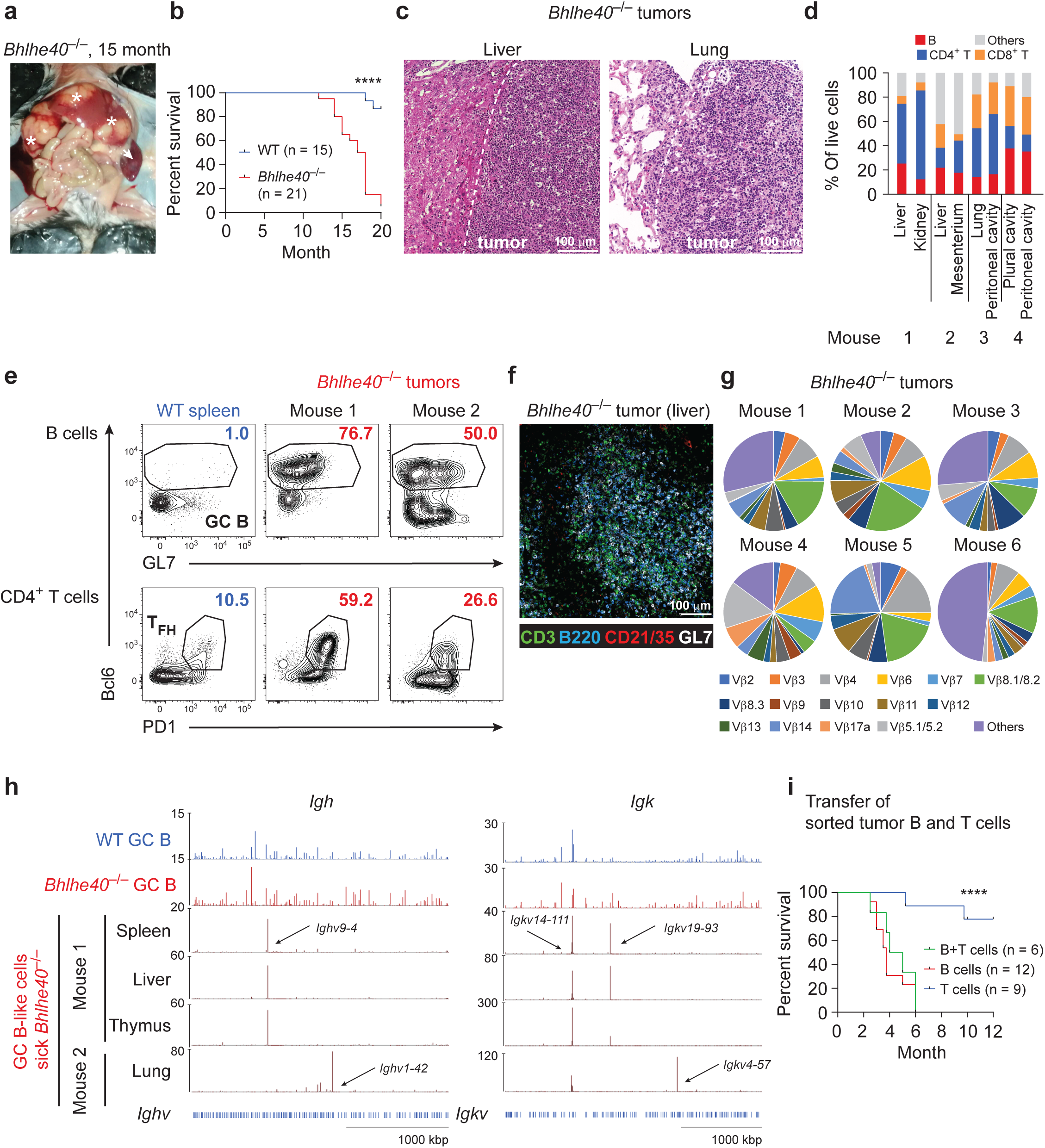
Lymphomagenesis in ageing Bhlhe40^−/−^ mice. (**a**) Image of a 15 month-old *Bhlhe40*^−/−^ mouse demonstrating splenomegaly (arrow) and nodular structures (asterisks) in the liver. (**b**) Kaplan-Meier survival curve depicting the survival of ageing WT and *Bhlhe40*^−/−^ mice. Pooled data for WT n = 15 and *Bhlhe40*^−/−^ n = 21 mice are shown. (**c**) H&E staining of sections of nodules in the liver and lung from two 15 month-old *Bhlhe40*^−/−^ mice. (**d**) Quantification of the composition of nodules of sick *Bhlhe40*^−/−^ mice as assessed by flow cytometry. (**e**) Flow cytometric analysis of GL7, PD1 and Bcl6 expression in the CD19^+^ B cell and CD4^+^ T cell population comparing an aged WT spleen and tumors from sick *Bhlhe40*^−/−^ mice between 15 and 18 month of age. Data are representative of n = 12 sick *Bhlhe40*^−/−^ mice. (**f**) Representative image of a tumor nodule section from the liver of a sick *Bhlhe40*^−/−^ mouse. CD3 (green), B220 (blue), CD21/CD35 (red) and GL7 (white) expression are shown. Data are representative of three analyzed mice. (**g**) TCRβ V segment usage by T_FH_-like cells from nodules of sick *Bhlhe40*^−/−^ mice as determined by flow cytometry. (**h**) Genome browser view of V segments in the *Igh* and *Igk* locus as determined by RNA-seq analysis of GC B cell-like cells of two sick *Bhlhe40*^−/−^ mice. (**i**) Kaplan-Meier survival curve depicting the survival of *Rag2*^−/−^ recipients that were transferred with sorted B and T cells from a sick *Bhlhe40*^−/−^ mouse. Pooled data from two independent experiments with n = 12 mice for B cells, n = 9 for T cells and n = 6 for T+B cells are shown.

## DISCUSSION

An efficient immune response to pathogens and vaccines relies on the interplay between two specialized lymphocyte subsets – GC B cells and T_FH_ cells – that takes place in the course of the GC reaction. Intriguingly, a number of transcriptional regulators have cell-intrinsic functions in both of these cell types. While many positive regulators of GC B cells and T_FH_ cells were identified to date, little is known about the negative regulation of the GC reaction. In this study, we identified the transcription factor Bhlhe40 as a novel common negative regulator of both B and T cell sides of the GC reaction. Regulation of two very different cell types involved in the same process by highly overlapping sets of transcription factors seems rather unusual and reasons for this overlap remain unclear. It is interesting to note that many of these shared transcription factors, including Bhlhe40, are rapidly upregulated upon BCR and TCR signaling and therefore seem to represent a conserved early response program downstream of both antigen receptors. As the GC reaction as well as the events preceding formation of GCs rely on recurring antigen receptor signaling events both in GC B cells and T_FH_ cells, such early response genes should be repeatedly induced in both subsets. Moreover, the complexity and highly dynamic nature of the GC reaction may allow to reveal relatively weak phenotypes that could be more difficult to capture in other settings – therefore resulting in the identification of factors with functions both in B and T cells even when these functions are subtle. Of note, most of the transcription factors shared by GC B and T_FH_ cells are not exclusively involved in regulation of only B and T cell subsets participating in the GC reaction. This applies to Bhlhe40 that regulates cytokine production in a variety of T cell subsets^42–45^ and, together with Bhlhe41, contributes to regulation of the development and self-renewal of B-1a B cells^38^. It is also conceivable that shared regulators of the GC reaction could in some cases represent a convenient ‘switch’ that would allow to enhance or dampen both B and T cell sides of the GC response by the same upstream signals. Indeed, one could envision that if Bhlhe40 could be up- or down-regulated both in B and T lymphocytes, for example by a common cytokine, this would allow simultaneous negative or positive regulation of both main lymphocyte types involved in the GC reaction.

Although Bhlhe40 recently emerged as an important regulator of cytokine production by T cells^42–45^, under the immunization conditions that favor T_FH_ cell differentiation used in our study, Bhlhe40- deficient antigen-specific CD4 T cells did not exhibit changes in expression of cytokine-encoding genes. We also did not observe dysregulation of expression of mitochondria-related genes observed in Bhlhe40- deficient tissue-resident memory CD8 T cells^41^. Instead, activated Bhlhe40-deficient CD4 T cells in our experiments upregulated cell cycle-related genes and exhibited increased proliferation. This is in line with several reports that together suggest that Bhlhe40 and Bhlhe41 may have a mild cytostatic function across a variety of cellular contexts^37^. As we also previously observed antiproliferative effects of these transcription factors in B-1a cells^38^, we initially hypothesized that Bhlhe40 also restrained proliferation of early GC B cells or their immediate precursors. However, extensive *in vivo* and *in vitro* experiments failed to reveal any evidence for a cytostatic function of Bhlhe40 in activated B cells. The earliest GC B cells also exhibited remarkably low levels of apoptosis and no change in frequency of apoptotic cells was observed between WT and KO cells. Moreover, while Bhlhe40 was expressed in highly proliferative common activated precursors of early MBCs, PBs, and GC B cells, only the later population was selectively affected by Bhlhe40 deficiency. Taken together, these results suggested that Bhlhe40 negatively regulates GC B cell development. In line with this notion, analysis of GC B cell differentiation trajectory by scRNA-seq indicated that Bhlhe40-deficient B cells gained a competitive advantage at the earliest steps of GC B cell differentiation. Moreover, Bhlhe40-deficient activated B cells exhibited a mild upregulation of GC B cell signature genes, suggesting that they may be more prone to adopt the GC B cell fate. Taken together, these results suggest that Bhlhe40 restrains the generation of the earliest cells seeding the GC reaction.

The negative regulation of early GC B cell generation, revealed through loss-of-function experiments in this study, would suggest that overexpression of Bhlhe40 should interfere with the GC reaction. Indeed, corroborating our results reported here, the transgenic overexpression of *Bhlhe40* driven by the VH promoter and the Eμ enhancer was previously shown to impair the generation of GC B cells upon immunization^48^.

Although *Bhlhe40*^−/−^ B cells did not exhibit altered levels of Bcl6 expression, the GC B cell phenotype of Bhlhe40 deficiency was highly reminiscent of that observed in I*μ*Bcl6 transgenic mice. Activated I*μ*Bcl6-transgenic B cells exhibited a 25% upregulation of Bcl6 protein over WT level already from the earliest stages of GC B cell differentiation^49^. This mild overexpression resulted in a competitive advantage of the transgenic cells selectively in the GC B cell compartment in the first days of the response but, very similarly to Bhlhe40 deficiency, did not affect fitness of the cells at later timepoints^49^. With age, I*μ*Bcl6 animals developed a lymphoma of GC B cell origin^50^.

Dysregulation of the GC reaction in Bhlhe40-deficient mice likewise culminated in the development of a GC B cell lymphoma. Cellular composition of these lymphomas strongly suggests that dysregulation of the GC reaction underlies the disease. However, the relative contribution of B cell- and T_FH_ cell-intrinsic aspects of the *Bhlhe40* KO phenotype to lymphomagenesis as well as possible involvement of the other cell types in the process remain to be tested. For example, it seems probable that the age-dependent decrease in regulatory T cells observed in Bhlhe40-deficient mice^32^ can also contribute to the lymphomagenesis.

Lymphomas in *Bhlhe40*^−/−^ mice took more than a year to develop, suggesting that accumulation of additional mutations in GC B cells in the course of a dysregulated GC reaction may be required for the malignant transformation. In line with this notion, RNA-seq analysis of the tumor cells demonstrated that they express *Aicda*, suggesting that lymphoma cells could continuously undergo AID-mediated mutagenesis. Expression of the *Igha* constant region by one of the two tumors subjected to RNA-seq suggests that the lymphoma may have originated from an IgA-switched cell from a ‘chronic’ GC in mucosal-associated lymphoid tissue. While it remains to be seen if Bhlhe40 mutations can be found in human B cell lymphomas, the unusual cellular composition of the lymphomas in Bhlhe40-deficient mice is reminiscent of certain human malignancies. The highly heterogeneous cellular composition of the nodules in diseased *Bhlhe40*^−/−^ mice contrasts with the dominance of transformed B cells in the case of human follicular lymphoma, GC B cell-like diffuse large B cell lymphoma and Burkitt’s lymphoma, and the normal GC B cell phenotype of the cells is inconsistent with classical Hodgkin’s lymphomas. However, the observed phenotype has intriguing similarities to human T cell/histiocyte-rich large B cell lymphoma and nodular lymphocyte-predominant Hodgkin’s lymphoma, as in both conditions transformed Bcl6-positive GC B cells are outnumbered by T cells^46, 47^. Of note, the etiology of these human lymphomas is not well understood, and it remains unknown what mutations lead to malignant transformation in these cases.

Taken together, the results reported in this study identify the transcription factor Bhlhe40 as a novel cell-intrinsic negative regulator of both B and T cell sides of the GC reaction, expression of which is crucial to prevent lymphomagenesis.

## MATERIAL AND METHODS

### Mice

All mice used in this study were maintained on the C57BL/6 genetic background. The *Igh*^B1-8hi 34^, *Bhlhe40*^−/− 31^, *Bhlhe41*^−/− 51^, *Rag2*^−/− 52^, *Mb1*-Cre ^53^, *Cd4*-Cre^54^, *Cd23*-Cre^55^, *Aicda*-Cre^55^ and *Vav*-Cre^56^ mice were described previously. *Bhlhe40*^fl/fl^ mice were generated as described below. WT C57BL/6J mice were obtained from Janvier Labs or bred in house. Mice analyzed in this study were at least 6 weeks old and kept under specific-pathogen-free conditions. Mice were bred and maintained at the Comparative Medicine Biomedicum facility of Karolinska Institutet (Stockholm, Sweden) or at the Research Institute for Molecular Pathology (Vienna, Austria). All mouse experiments were carried out according to valid project licenses, which were approved and regularly controlled by the Swedish and/or Austrian Veterinary Authorities.

*Generation of Bhlhe40*^fl/fl^ *mice*

*Bhlhe41*^fl^ mice were generated by CRISPR–Cas9-mediated genome editing in mouse zygotes^57^. A solution containing two sgRNAs targeting *Bhlhe40* sequences in introns 2 and 4 (Supplementary Fig. 6a), Cas9 mRNA, Cas9 protein and ssDNA repair template was injected into C57BL/6 x CBA zygotes. Correct targeting was confirmed by sequencing and the mice were backcrossed to the C57BL/6 genetic background for 4 generations before crossing them to the Cre lines.

### Flow cytometry

Mouse organs were harvested, and single cell suspensions were obtained by mincing through 70 μm cell strainers. For detection of antigen-specific CD4 T cells, splenic single cell suspensions were incubated with APC-conjugated I-A^b^ Mtb Ag85b_280-294_ and I-A^b^ 2W1S tetramers (provided by National Institutes of Health tetramer core facility) for 1 hour prior to Fc receptor blocking and further cell surface staining.

For detection of PE-specific B cells in the polyclonal response in mice, splenic single cell suspensions were incubated in IMDM (Thermo Fisher Scientific) pH 3.0 on ice for 1 min to remove Fc receptor-bound antibodies, washed twice with PBS/2% FCS and stained with PE (Prozyme, 20 µg/ml). RNA flow cytometry measurement of *Bhlhe40* expression was performed using the PrimeFlow RNA Assay (Thermo Fisher Scientific) according to the manufacturer’s instructions. High-sensitivity Alexa Fluor 647 probes were used. Intracellular staining for cleaved caspase-3 was performed using the 5A1E rabbit monoclonal antibody, Alexa Fluor 647–labeled anti–rabbit F(ab)_2_ fragment (both from Cell Signaling Technology) and the Cytofix/Cytoperm kit (BD) according to manufacturer’s instructions. Intracellular staining for transcription factors was performed using the Foxp3 staining buffer set (Thermo Fisher Scientific) according to manufacturer’s instructions. Detection of cells positive for 5-Ethynyl-2′- deoxyuridine (EdU) was performed using the Click-iT Plus EdU A488 or the Click-iT Plus Pacific Blue Flow Cytometry Assay Kit (Thermo Fisher Scientific) following manufacturer’s instructions with minor modifications. Data were acquired on an LSR Fortessa Flow Cytometer (BD Biosciences) and analyzed with FlowJo software v.10 (BD).

### Flow cytometry antibodies

For staining of murine samples, monoclonal antibodies specific for BCL6 (REA373 and K112-91), CD4 (GK1.5, RM4-8, S11), CD3e (145-2C11, 17A2, REA641), CD8 (53-6.7), PD-1 (RMP1-30, REA802, 29F.1A12), CXCR5 (2G8, REA215), CD44 (IM7), CD62L (MEL-14), CD19 (1D3, 6D5), B220 (RA3-6B2), GL7 (GL7), CD95 (JO2 and REA453), CCR6 (REA277 and 29-2L17), CD38 (90), CD138 (REA104 and 281-2), TACI (8F10), CD86 (GL1), CXCR4 (2B11), IgM (REA979 and RMM-1), IgD (11-26c2a), IgG1 (REA1017 and RMG1-1), IgL (R26-46 and JC5-1), CD21/CD35 (7E9), CD45.1 (A20, REA1179), CD45.2 (104, 104-2), IgK (187.1), Gr-1 (RB6-8C5), F4/80 (BM8), CD11c (HL3), NK1.1 (PK136), TCRβ (H57-597, REA318), TCRγδ (GL3), and Ter119 (Ter119) were purchased from BioLegend, BD Biosciences, Miltenyi Biotec or Thermo Fisher Scientific and were used at dilutions specified by the manufacturer or determined experimentally. For staining for TCRβ subtypes in *Bhlhe40*^−/−^ tumor mice, the mouse Vβ TCR screening panel (BD bioscience) was used.

### Generation of mixed BM chimeras and adoptive cell transfers

BM cells from WT and *Bhlhe40*^−/−^, *Bhlhe41*^−/−^, or *Bhlhe40*^−/−^*Bhlhe41*^−/−^ mice were stained with CD4, CD8, CD3, TCRβ, TCRγδ, NK1.1, CD19, CD11b, CD11c, Gr-1, and Ter119 APC- or PE-labeled antibodies followed by magnetic depletion with anti-APC or -PE Micro-Beads (Miltenyi Biotec), respectively. WT and KO cells were mixed at a 1:1 ratio (unless stated otherwise) and 2-4 x 10^6^ cells were transferred intravenously into lethally irradiated (split dose – 500 rads, twice) WT or *Rag2*^−/−^ recipients (as indicated). CD45.1 and CD45.2 congenic markers were used as indicated. Chimeras were analyzed > 6 weeks after reconstitution. For adoptive transfer experiments, splenocytes from congenically distinguishable *Bhlhe40*^+/+^ *Igh*^B1-8hi/+^ and *Bhlhe40*^−/−^ *Igh*^B1-8hi/+^ mice were mixed targeting a 1:1 ratio of WT and *Bhlhe40*^−/−^ cells in the antigen-specific B cell compartment (CD19^+^Igλ^+^). The ratio was confirmed by flow cytometric analysis of an aliquot from the mix and adjusted when necessary. 10-14 x 10^6^ of splenocytes (∼3-5 x 10^5^ NP-specific B cells counting both donors together) were injected into the tail vein of congenically distinguishable sex-matched C57BL/6 recipient mice. Recipient mice were immunized 12-24 hours after transfer as described below.

For adoptive transfer of B and T cells from *Bhlhe40*^−/−^ tumor mice, CD19^+^ B cells and CD4^+^ T cells were sorted from the spleens of sick *Bhlhe40*^−/−^ tumor mice using a Sony SH800 Cell Sorter. >5 x 10^5^ of B and/or T cells were transferred intravenously into congenically distinguishable sex-matched *Rag2*^−/−^ recipient mice via injection into the tail vein.

### Immunizations experiments

For B1-8^hi^ transfer experiments, C57BL/6 recipient mice were pre-immunized by intraperitoneal injection of 100 µg OVA (BioSearch Technologies) dissolved in PBS and precipitated in alum (Sigma-Aldrich) at a 1:1 ratio > 2 weeks prior to adoptive cell transfer. 12 to 24 hours after B1-8^hi^ cell transfer, recipient mice were immunized by intraperitoneal injection of 100 µg NP19-OVA (BioSearch Technologies) precipitated in alum (Sigma-Aldrich) at a 1:1 ratio. For *in vivo* proliferation analysis, 1 mg 5-Ethynyl-2′-deoxyuridine (EdU, Sigma-Aldrich) diluted in PBS was intraperitoneally or intravenously injected at indicated time-points post immunization. For Phycoerythrin (PE) immunization, mice were subcutaneously injected with 50 μl of an emulsion containing 50 μg of PE. Emulsions were prepared by mixing PE in PBS in a 1:1 ratio with Complete Freund’s Adjuvant (Sigma-Aldrich). For peptide immunizations, mice were intraperitoneally injected with 100 µg Ag85b_280-294_ (FQDAYNAAGGHNAVF) or 2W1S (EAWGALANWAVDSA) peptide in 100 µl of a 1:1 emulsion with Complete Freund’s adjuvant (Sigma-Aldrich).

### Quantitative RT-PCR (qRT-PCR)

WT splenic naïve B cells were isolated using CD43 Micro-Beads (Miltenyi Biotec) and stimulated with anti- CD40 (1 µg/ml, Invitrogen) or anti-IgM (1 µg/ml, Jackson ImmunoResearch) antibody alone or in combination with IL-4 (20 ng/ml, Peprotech) for up to 1 day. Total RNA was isolated from cultured B cells at indicated timepoints using the Quick-RNA Microprep Kit (Zymo Research). cDNA was synthesized with the RevertAid RT Kit (Thermo Scientific) and qRT-PCR was performed on a CFX96-C1000 Thermo Cycler (Bio-Rad) using the SensiFAST SYBR No-ROX Kit (Bioline, Meridian Bioscience). *Hprt1* was used as a housekeeping gene and the standard curve method was used for quantification. The following primers were used: Bhlhe40-F CTC CTA CCC GAA CAT CTC AAA C, Bhlhe40-R CCA GAA CCA CTG CTT TTT CC, Hprt1 -F AGT GTT GGA TAC AGG CCA GAC, Hprt1-R CGT GAT TCA AAT CCC TGA AGT.

### H&E stainings

Tissue samples of liver and lung from *Bhlhe40*^−/−^ tumor mice were processed using the standard tissue protocol on Automatic Tissue Processor Donatello (Diapath) and paraffin-embedded. Then 2 µm paraffin sections were cut. Sections were rehydrated with xylene substitute (Thermo Scientific) and ethanol, followed by staining with hematoxylin and eosin (both Thermo Scientific) for 1 min each and again dehydrated with ethanol and xylene substitute. After embedding in aqueous (Eukitt Neo) mounting medium, slides were analyzed using a Pannoramic FLASH 250 III slide scanner equipped with an Adimec Quartz Q12A180 camera.

### Immunofluorescence microscopy

For confocal immunofluorescence microscopy, organs were harvested, fixed in 4% PFA in PBS for 1 to 2 hours and subsequently incubated in 30% sucrose in PBS for up to 3 hours. Organs were embedded in Tissue Tek Optimal Tissue Cutting Temperature medium (OCT, Sakura Finetek) and snap frozen on dry ice. 8 µm sections were cut using a cryostat and sections were mounted on SuperFrost Plus glass slides (ThermoFisher Scientific). Sections were incubated in ice-cold acetone for 10 min prior to blocking in 5% BSA in PBS. To block endogenous biotin, the streptavidin/biotin blocking kit (Vector Laboratories) was used according to manufacturer’s instructions. Staining was performed for 1 hour at RT and slides were mounted with Fluoromount Aqueous Mounting Medium (Sigma-Aldrich). Confocal images were acquired on an LSM 700 system (Carl Zeiss) with 405-, 488-, 555-, and 639 nm excitation lines at the Biomedicum Imaging Core at the Karolinska Institute. Images were processed using the ZEN 2.3 Black Edition (Carl Zeiss) or Imaris (Bitplane) Imaging software.

To analyze the size of the GCs in spleen sections of untreated WT and *Bhlhe40*^−/−^ mice, samples were prepared and imaged as described above. Additional images were acquired using a Pannoramic 250 Flash Slide Scanner equipped with a x20 objective (3DHISTECH). GCs were identified as IgD-negative areas within B cell follicles and the CaseViewer (3DHISTECH) or ImageJ software were used to measure the size of the GCs.

### ChIP-seq analysis of Bhlhe40 binding

WT splenic naïve B cells were isolated using CD43 Micro-Beads (Miltenyi Biotec) and stimulated with 1 µg/ml anti-CD40 antibody (Invitrogen) for 4 hours. Cells were fixed with 1% PFA in PBS in the presence of 1% FCS at room temperature. Fixation was stopped after 10 minutes by addition of glycine in PBS (final concentration 0.1 M). Cells were washed with 0.1 M glycine solution in PBS and pellets were frozen. ChIP and library preparation was performed as recently described^58^ with minor modifications. 3x10^6^ and 22x10^6^ fixed frozen B cells were thawed at room temperature and diluted with SDS lysis buffer (50mM Tris/HCl pH8, 0.5% SDS and 10mM EDTA pH8) containing 1x cOmplete protease inhibitor. Cells were sonicated in a Bioruptor Plus sonicator (Diagenode). To neutralize the SDS, Triton X100 was added to a final concentration of 1% along with cOmplete protease inhibitor. Samples were incubated at room temperature for 10 minutes. 3 μg of rabbit polyclonal anti-Bhlhe40 antibody (Novus Biologicals, NB100- 1800, lot C2) was added to 10 μl Protein G-coupled Dynabeads (Thermo Fisher Scientific) in PBS with 0.5% BSA and incubated with rotation for 4 hours at 4°C. Antibody-coated Dynabeads were washed with PBS with 0.5% FCS and mixed with cell lysate in PCR tubes. Tubes were incubated rotating overnight at 4°C. Immunoprecipitated chromatin was washed with 150 μl of low-salt buffer (50 mM Tris/HCl, 150 mM NaCl, 0.1% SDS, 0.1% Sodium Deoxycholate, 1% Triton X-100, and 1 mM EDTA), high-salt buffer (50 mM Tris/HCl, 500 mM NaCl, 0.1% SDS, 0.1% Sodium Deoxycholate, 1% Triton X-100, and 1 mM EDTA) and LiCl buffer (10 mM Tris/HCl, 250 mM LiCl, 0.5% IGEPAL CA-630, 0.5% Sodium Deoxycholate, and 1 mM EDTA), followed by two washes with TE buffer (10 mM Tris/HCl and 1 mM EDTA) and two washes with ice-cold Tris/HCl pH 8. For tagmentation, bead bound chromatin was resuspended in 30 μl of tagmentation buffer. 1 μl of transposase (Nextera, Illumina) was added and samples were incubated at 37°C for 10 minutes followed by two washes with low-salt buffer. Bead bound tagmented chromatin was diluted in 20 μl of water. 25 μl PCR master mix (Nextera, Illumina) and 5 μl indexed amplification primers^59^ (0.125 μM final concentration) were added and libraries prepared using the following PCR program: 72°C 5 minutes (adapter extension); 95°C 5 minutes; followed by 11 cycles of 98°C 10 seconds, 63°C 30 seconds and 72°C 3 minutes. All steps from sonication to library amplification PCR were performed in the same PCR tube. After PCR amplification, library cleanup was done using Agencourt AmPureXP beads (Beckman Coulter) at a PCR mix to bead ratio of 1:1. DNA concentrations in purified samples were measured using the Qubit dsDNA HS Kit (Invitrogen). Libraries were subjected to Illumina deep sequencing (NextSeq 500).

### Analysis of ChIP-seq data

Bhlhe40 ChIP-seq reads from our activated B cell dataset (3x10^6^ and 22x10^6^ B cells) as well as a published activated CD4 T cell ChIP-seq dataset (GSE113054 in GEO database)^42^ were aligned to the mouse genome assembly version of July 2007 (NCBI37/mm9), using the Bowtie2 program ^60^ (usegalaxy.eu; galaxy version 2.3.4.2 ^61^) and bam files for 3x10^6^ and 22x10^6^ cell samples were merged. Peaks were called with a *P* value of < 10^−10^ by using the MACS program version 1.3.6.1 ^62^ with default parameters. The identified peaks were then assigned to target genes as described^63^. Bhlhe40 peaks overlapping with the transcription start site (TSS) were referred to as promoter peaks. For motif discovery, we used the MEME-ChIP suite (usegalaxy.eu; galaxy version 4.11.2^61^)^64^ to predict the most significant motifs present in the 300 bp centered at the peak summit of the top 500 sequences, as sorted by the fold enrichment score of the MACS program.

### RNA-seq analysis

For RNA-seq analysis of activated B cell populations, congenically distinguishable WT and KO B1-8^hi^ B cells were co-transferred into OVA-preimmunized WT recipients as described above and isolated at day 4 after NP-OVA immunization. Splenocytes were depleted using Igκ/CD4/CD8/GR1/Ter119/TCRb/TCRgd/CD11b/CD11c/NK1.1 antibodies and anti-PE Micro-Beads (Miltenyi Biotec), and WT (CD45.1^+^CD45.2^−^) and KO (CD45.1^+^CD45.2^+^) antigen-specific B cells (CD19^+^Igλ^+^) GC B cells (CCR6^−^GL7^hi^) or non-GC B activated B cells (CCR6^+^GL7^int^) were double sorted. For the second sort, 500 cells were sorted directly into lysis buffer and subjected to the SMART-Seq v2 Cell protocol^65^. Libraries were subjected to Illumina deep sequencing (HiSeqV4 SR50).

For RNA-seq analysis of activated T cells, WT and *Bhlhe40*^−/−^ mixed BM chimeras were generated as described above and intraperitoneally injected with 100 µg Ag85b280-294 peptide in 100 µl of a 1:1 emulsion with Complete Freund’s adjuvant (Sigma-Aldrich). On day 11 post immunization, spleens were harvested and single cell suspensions were incubated with APC-conjugated I-A^b^ Mtb Ag85b_280-294_ tetramer (provided by National Institutes of Health tetramer core facility) for 1 hour at 4°C and tetramer-positive cells were enriched using anti-APC Micro-Beads (Miltenyi Biotec). CD19^−^CD11c^−^CD11b^−^F4/80^−^CD8a^−^ WT (CD45.1^+^CD45.2^+^) and *Bhlhe40*^−/−^ (CD45.1^+^) antigen-specific T cells (Tetramer^+^CD4^+^TCRb^+^) were double sorted. For the second sort, 1000 cells were directly sorted into single cell lysis solution (Invitrogen) according to manufacturer’s instructions. cDNA and second strand synthesis was performed using NEBNext Ultra II RNA First Strand Synthesis (NEB cat E7771) and NEBNext Ultra II Directional RNA Second Strand Synthesis (NEB Cat E7550) modules. QIAseq FastSelect rRNA HMR kit (Qiagen) was utilized to minimize rRNA contribution to the final library. Custom made Tn5 (transposase) in combination with oligo replacement and PCR amplification was utilized to generate indexed sequencing libraries^66^. Libraries were pooled and paired-end sequenced (41 cycles) using the NextSeq 500 system (Illumina).

Tumor B cells from indicated organs were sorted as CD19^+^CD95^+^GL7^+^ (mouse 1) or CD19^+^CD95^+^ (mouse 2). Libraries were prepared as previously described ^38^ with minor modifications. In brief, RNA was isolated with an RNeasy Plus Mini or Micro Kits (Qiagen), and mRNA was obtained by poly(A) selection with a Dynabeads mRNA purification kit (Thermo Fisher Scientific), followed by fragmentation by heating at 94 °C for 3 min (in fragmentation buffer). The fragmented mRNA was used as a template for first-strand cDNA synthesis with random hexamers and a Superscript VILO cDNA Synthesis kit (Thermo Fisher Scientific). The second-strand cDNA was synthesized with 100 mM dATP, dCTP, dGTP and dUTP in the presence of RNase H, E. coli DNA polymerase I and DNA ligase (Thermo Fisher Scientific). Sequencing libraries were prepared with the NEBNext Ultra II DNA Library Prep Kit for Illumina (New England BioLabs). For strand-specific RNA-sequencing, the uridines present in one cDNA strand were digested with uracil-N-glycosylase (New England BioLabs), as described ^67^, followed by PCR amplification with NEBNext UltraII Q5 Master Mix (New England BioLabs). Libraries were subjected to Illumina deep sequencing (HiSeqV4 SR50). For RNA-seq analysis of polyclonal WT and *Bhlhe40*^−/−^ GC B cells LZ (CD19^+^CD95^+^GL7^+^CXCR4^lo^CD86^hi^) and DZ (CD19^+^CD95^+^GL7^+^CXCR4^hi^CD86^lo^) GC B cells were sorted from non-immunized mixed BM chimeras and subjected to RNA-seq as discussed above for tumor cells. Tracks from DZ GC B cells are shown in Figure 6h.

### Bioinformatic analysis of RNA-seq data

Sequence reads that passed the Illumina quality filtering were considered for alignment. Reads corresponding to mouse ribosomal RNAs (BK000964.1 and NR046144.1) were removed. The remaining reads were cut down to a read length of 44 bp and aligned to the mouse transcriptome (genome assembly version of July 2007 NCBI37/mm9) using TopHat version 1.4.1 ^68^. The calculation of RNA expression values was all based on the RefSeq database, which was downloaded from UCSC on January 10th, 2014. The annotation of immunoglobulin and T cell receptor genes was incorporated from the Ensembl release 67^69^. Genes with overlapping exons were flagged and double entries (i.e. exactly the same gene at two different genomic locations) were renamed. Genes with several transcripts were merged to consensus- genes consisting of a union of all underlying exons using the fuge software (I. Tamir, unpublished), which resulted in 25,726 gene models.

For analysis of differential gene expression, the number of reads per gene was counted using HTseq version 0.5.3 ^70^ with the overlap resolution mode set to ’union’. The data sets were analyzed using the R package DESeq2 version 1.2.10 ^71^. Sample normalizations and dispersion estimations were conducted using the default DESeq2 settings. Transcripts per million (TPM) were calculated from RNA-seq data, as described ^72^.

GSEA was performed using the GSEA software from the Broad Institute ^73^. Genes were ranked on the basis of their change in expression (log_2_ fold values) as determined by the DESeq2 package and were compared to gene sets from the MSigDB or defined gene sets from literature or our data. GC B cell signature was used from Mabbott et al ^39^ with *Bcl6* and *Gcsam* manually added to the signature. Early GC B cell signature was generated by pairwise comparison of our WT day 4 GC B and CCR6^+^ B cell datasets, using top 500 GC B cell-upregulated genes.

### scRNA-seq library preparation

For single cell sequencing, WT and *Bhlhe40*^−/−^ B1-8^hi/+^ B cells were co-transferred and sorted at day 3.5 (5 mice) and day 4 (5 mice) after immunization as described above for bulk RNA-seq. Cells from day 3.5 and 4 after immunization were pooled. An additional WT sample was sorted at day 2.5 after immunization. Lineage depletion for CD4/TCRβ/TCRγδ/CD11c/Gr1/NK1.1/Ter119/Igκ was performed using MACS anti-PE Micr-beads (Miltenyi Biotec). CD19^+^Igλ^+^CD45.1^+^CD45.2^−^ (WT) and CD19^+^Igλ^+^CD45.1^+^CD45.2^+^ (*Bhlhe40*^−/−^) cells were double sorted using a FACS Aria III sorter. Broad gates were applied to include PBs that may start downregulating some of these surface molecules. For day 3.5-4 samples, cells were processed using the 10X Genomics kit version 2 immediately after sorting. For day 2.5, cells were resuspended in cold PBS/0.04% BSA and fixed by adding 4 volumes 100% Methanol (VWR) and incubation for 30 min at -20 °C. Day 2.5 cells were stored at -80°C and also processed as described ^74^ using the 10X Genomics kit version 3. Cells were sequenced paired end with 75 bp read length on a NextSeq550 system (Illumina).

### Analysis of single cell RNA-seq data

The sequenced 10X Chromium libraries from two samples were mapped to mm10 mouse genome and assigned to droplets with Cell Ranger software (version 3.0.1) with default parameters ^75^. Transcriptomes of 7,324 WT cells with the median read coverage of 2,292 (days 2.5), 2,089 WT cells with a median read coverage of 2,507 (day 3.5-4) and 1,677 KO cells with a median read coverage of 3823 (day 3.5-4) were. Joined embedding for days 2.5 and 3.5-4 was performed with scanpy based on spliced transcripts from velocyto (version 0.17.13)^76^. First, the data were normalized and filtered with scanpy.pp.recipe_seurat workflow^77^. Next, UMAP embedding was performed with default parameters. Cells were clustered with louvain algorithm^78^ and cell populations were defined as groups of clusters identified by this algorithm. GC B cell signature was used from Mabbott et al^39^ with *Bcl6* and *Gcsam* manually added to the signature.

### Statistical analysis

Statistical analysis was performed with the GraphPad Prism 8 software. Two-tailed unpaired Student’s *t*-test analysis was used to assess the statistical significance of one observed parameter between two experimental groups. Two-tailed paired Student’s *t*-test analysis was used to assess the statistical significance of one observed parameter between two related groups. The statistical evaluation of the RNA-seq data is described above.

## ACKNOWLEDGEMENT

We thank G. Schmauß, M. Weninger and their colleagues for sorting by flow cytometry; A. Sommer’s team at the Vienna Biocenter Support Facilities GmbH (VBCF) for Illumina sequencing; A. Kavirayani and his team for histological service; Josefine Dunst for critical reading of the manuscript. This study was supported by the Swedish Research Council (grant 2017-01118 to T.K.), Cancerfonden (grant CAN 2018/710 to T.K.), Åke Wibergs Stiftelse (grant M18-0094 to T.K.), the Austrian Science Fund (grant P28841 to T.K.), and a stipend from the Wenner-Gren Foundations (to T.K.), a stipend from the German Research Foundation DFG (RE 4264/1-1 to A.R.), Boehringer Ingelheim (M.B. and R.R.) and the European Research Council (ERC) under the European Union’s Horizon 2020 research and innovation program (grant agreement 740349-PlasmaCellControl, to M.B.). We thank the National Institutes of Health tetramer core facility for the preparation of MHC tetramers.

## Notes

### Competing Interest Statement

The authors have declared no competing interest.

