## Supplementary material for "Cell-intrinsic functions of the transcription factor Bhlhe40 in activated B cells and T follicular helper cells restrain the germinal center reaction and prevent lymphomagenesis"

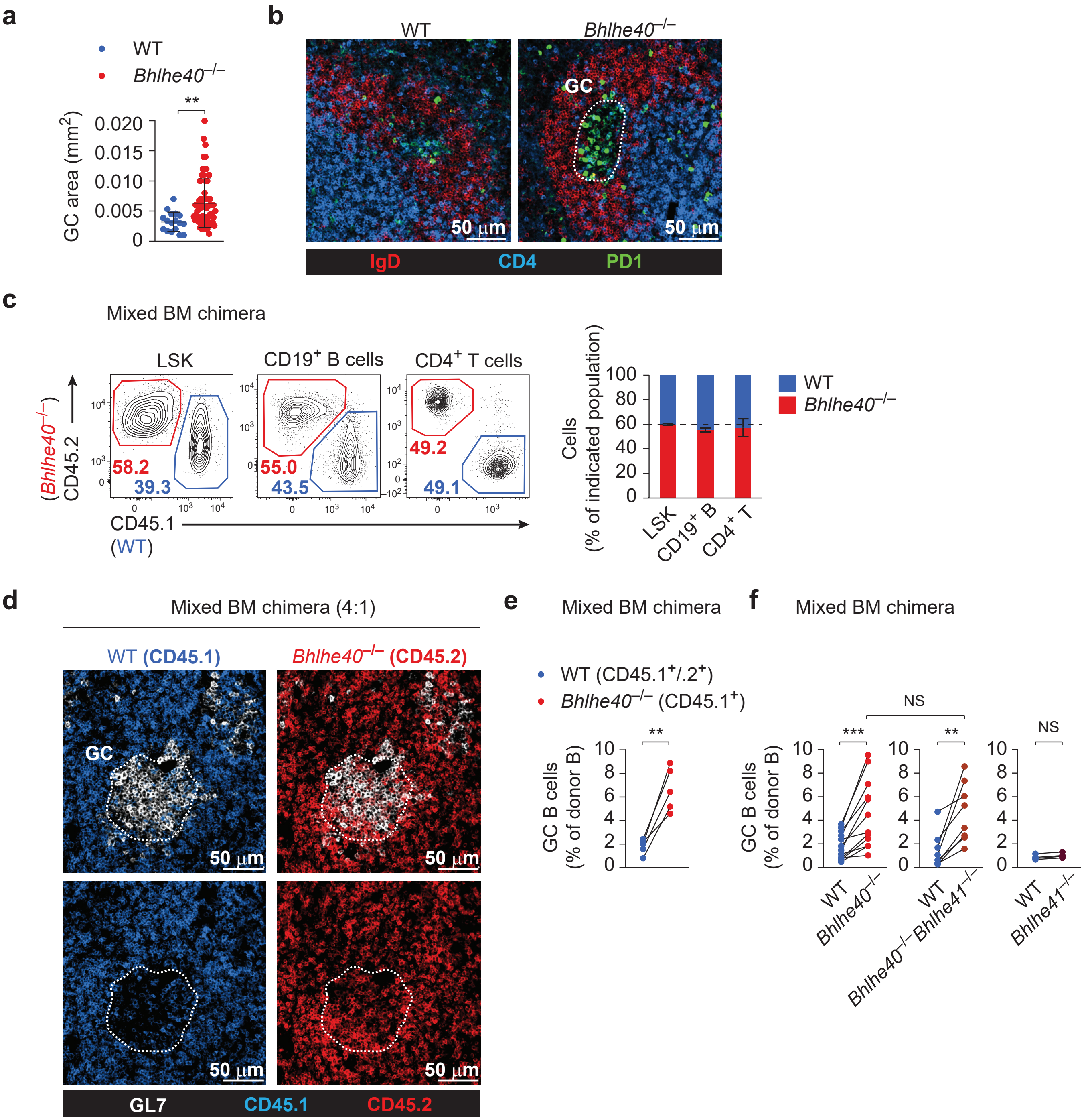
a
WT
Bhlhe40−/−
**
GC area (mm2)
0.020
0.015
0.010
0.005
0
b
WT
Bhlhe40−/−
GC
50 µm
50 µm
IgD
CD4
PD1
c
Mixed BM chimera
LSK
CD19+ B cells
CD4+ T cells
(Bhlhe40−/−) CD45.2
58.2
39.3
55.0
43.5
49.2
49.1
CD45.1
(WT)
Cells (% of indicated population)
100
80
60
40
20
0
LSK
CD19+ B
CD4+ T
WT
Bhlhe40−/−
d
Mixed BM chimera (4:1)
WT (CD45.1)
Bhlhe40−/− (CD45.2)
GC
50 µm
GL7
CD45.1
CD45.2
e
Mixed BM chimera
WT (CD45.1+/.2+)
Bhlhe40−/− (CD45.1+)
**
GC B cells (% of donor B)
f
Mixed BM chimera
NS
***
**
NS
WT
Bhlhe40−/−
WT
Bhlhe40−/− Bhlhe41−/−
WT
Bhlhe41−/−

Figure S1

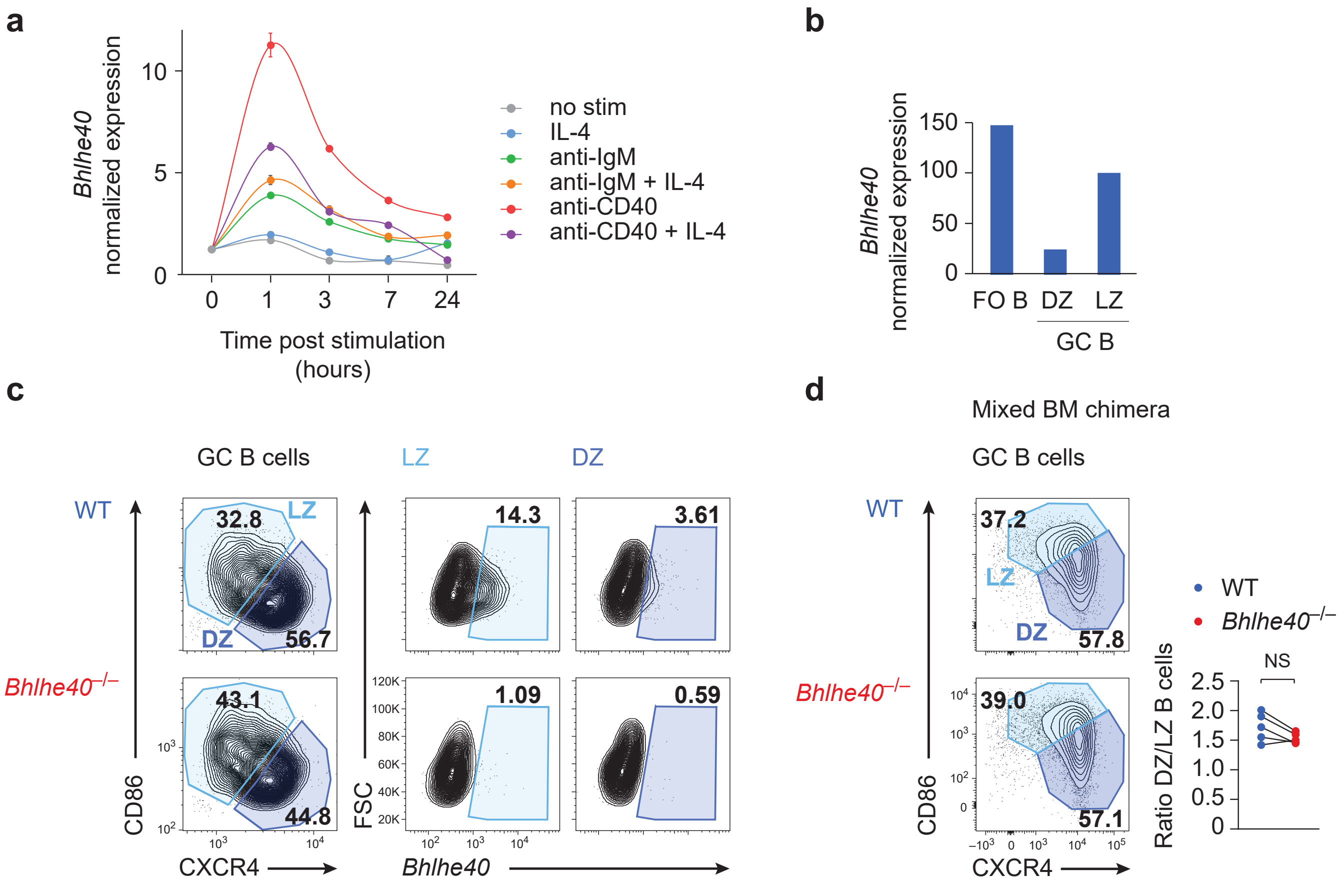

a
Bhlhe40 normalized expression
Time post stimulation (hours)
no stim
IL-4
anti-IgM
anti-IgM + IL-4
anti-CD40
anti-CD40 + IL-4
b
Bhlhe40 normalized expression
FO B
DZ
LZ
GC B
c
GC B cells
LZ
DZ
WT
Bhlhe40−/−
32.8
56.7
43.1
44.8
14.3
3.61
1.09
0.59
CD86
CXCR4
FSC
Bhlhe40
d
Mixed BM chimera
GC B cells
37.2
57.8
39.0
57.1
WT
Bhlhe40−/−
NS
Ratio DZ/LZ B cells


Figure S2

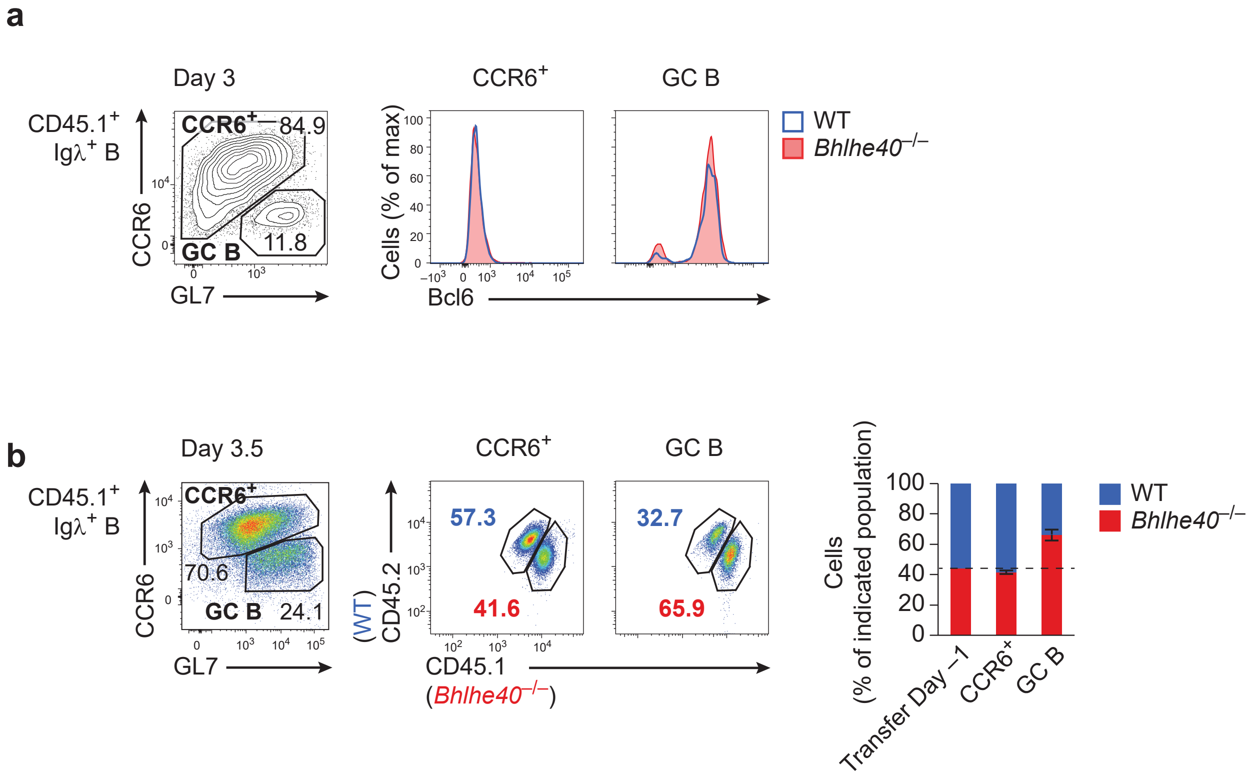

a
Day 3
CD45.1+
Igλ+ B
CCR6
GL7
CCR6+
84.9
GC B
11.8
CCR6+
GC B
Cells (% of max)
Bcl6
WT
Bhlhe40–/–
b
Day 3.5
CD45.1+
Igλ+ B
CCR6
GL7
CCR6+
70.6
GC B
24.1
CCR6+
GC B
57.3
41.6
32.7
65.9
(WT)
CD45.2
CD45.1
(Bhlhe40–/–)
Cells
(% of indicated population)
Transfer Day –1
CCR6+
GC B
WT
Bhlhe40–/–


Figure S3

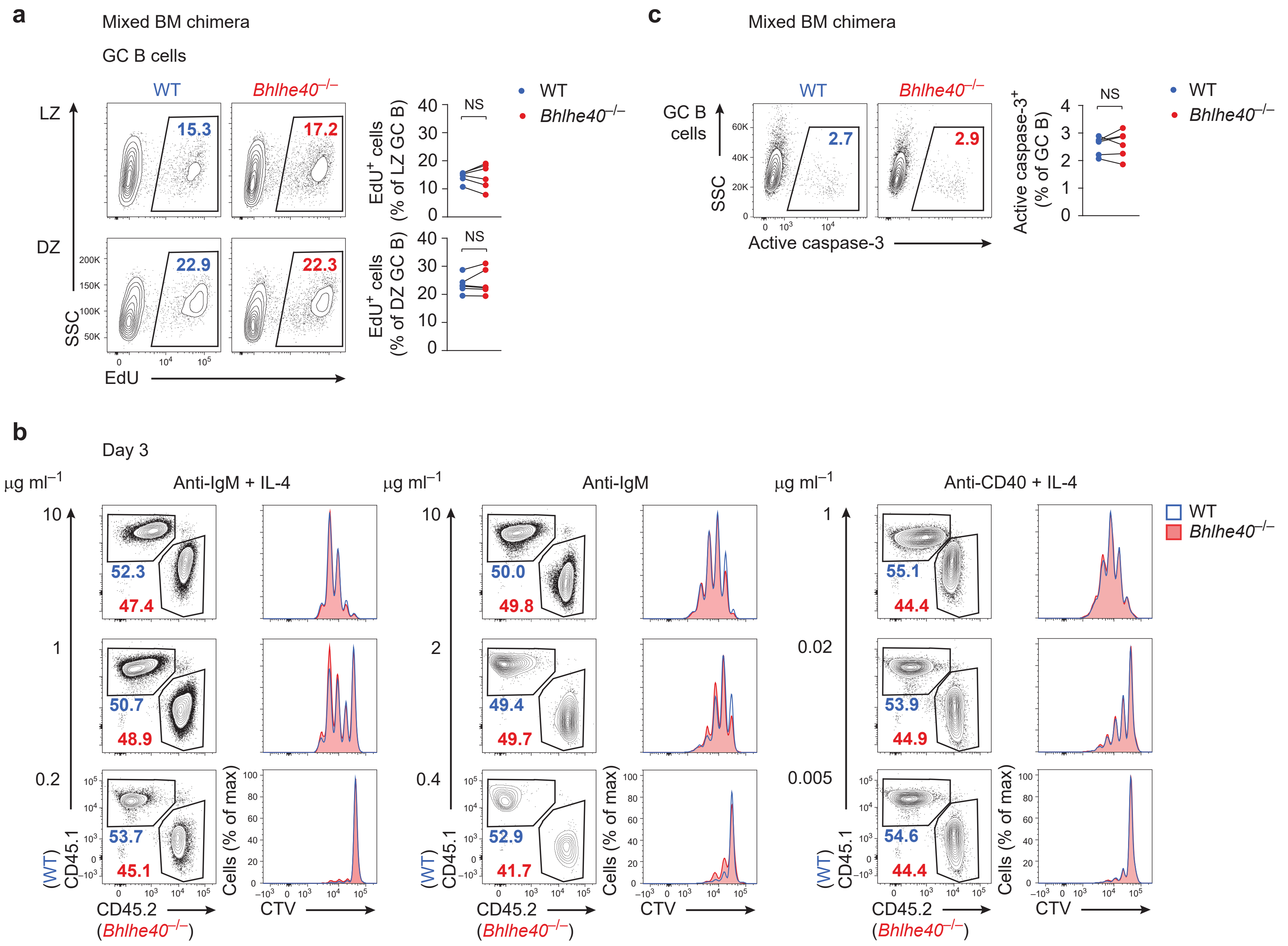
a
Mixed BM chimera
GC B cells
WT
Bhlhe40−/−
LZ
DZ
15.3
17.2
22.9
22.3
SSC
EdU
EdU+ cells (% of LZ GC B)
EdU+ cells (% of DZ GC B)
NS
c
Mixed BM chimera
GC B cells
2.7
2.9
Active caspase-3
Active caspase-3+ (% of GC B)
b
Day 3
μg ml−1
Anti-IgM + IL-4
Anti-IgM
Anti-CD40 + IL-4
52.3
47.4
50.7
48.9
53.7
45.1
50.0
49.8
49.4
49.7
52.9
41.7
55.1
44.4
53.9
44.9
54.6
44.4
CD45.1 (WT)
CD45.2 (Bhlhe40−/−)
Cells (% of max)
CTV

Figure S4

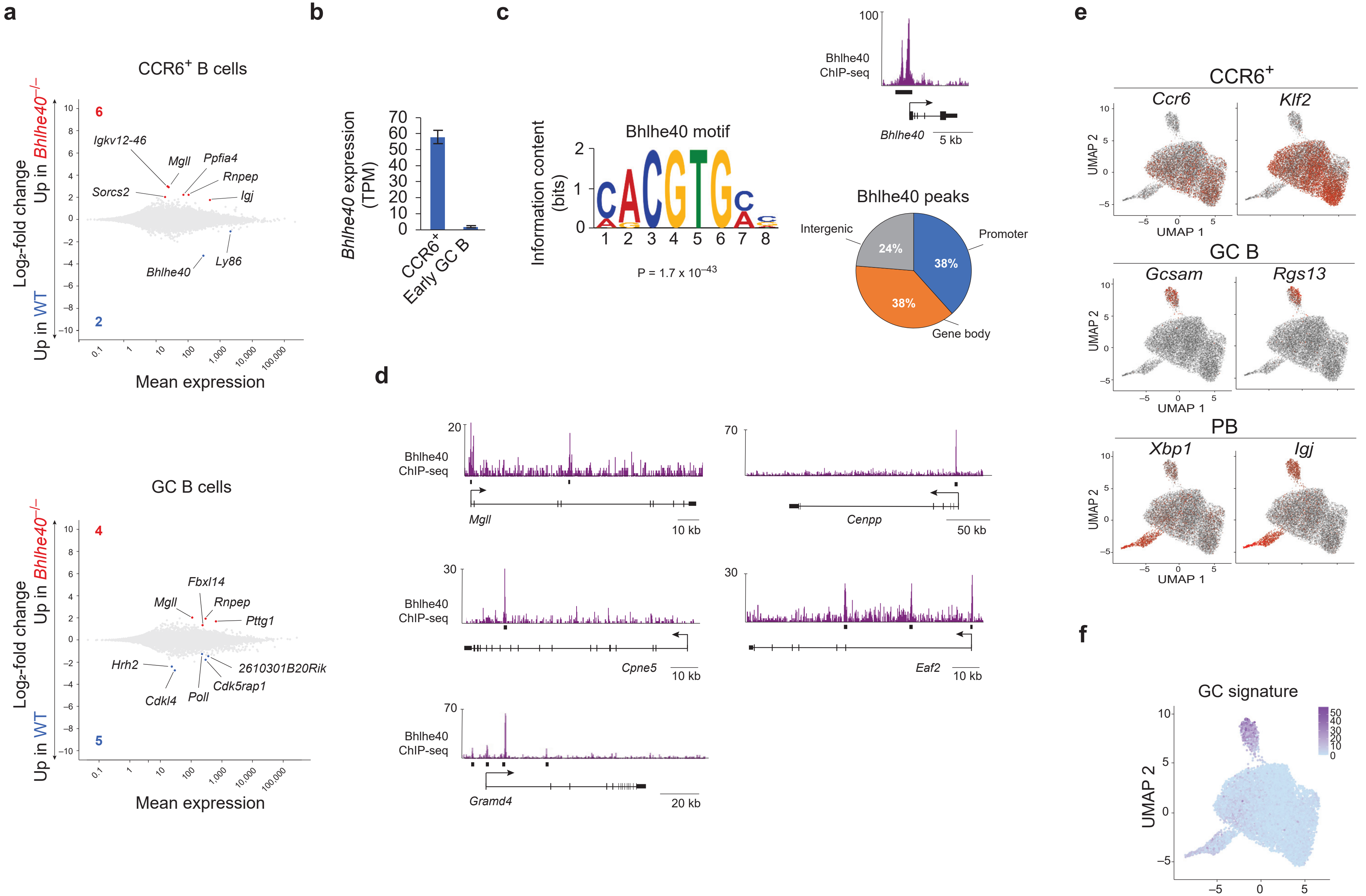
a
CCR6+ B cells
Up in Bhlhe40−/−
Log₂-fold change
Up in WT
6
2
Igkv12-46
Mgll
Ppfia4
Rnpep
Sorcs2
Igj
Ly86
Bhlhe40
Mean expression
GC B cells
4
5
Fbxl14
Mgll
Rnpep
Pttg1
Hrh2
2610301B20Rik
Cdk5rap1
Cdkl4
Poll
b
Bhlhe40 expression (TPM)
CCR6+
Early GC B
c
Bhlhe40 motif
Information content (bits)
P = 1.7 x 10−43
Bhlhe40 ChIP-seq
Bhlhe40
5 kb
Bhlhe40 peaks
Intergenic
Promoter
Gene body
24%
38%
38%
d
Bhlhe40 ChIP-seq
Mgll
10 kb
Cenpp
50 kb
Cpne5
10 kb
Eaf2
10 kb
Gramd4
20 kb
e
CCR6+
Ccr6
Klf2
GC B
Gcsam
Rgs13
PB
Xbp1
Igj
UMAP 1
UMAP 2
f
GC signature
UMAP 1
UMAP 2

Figure S5

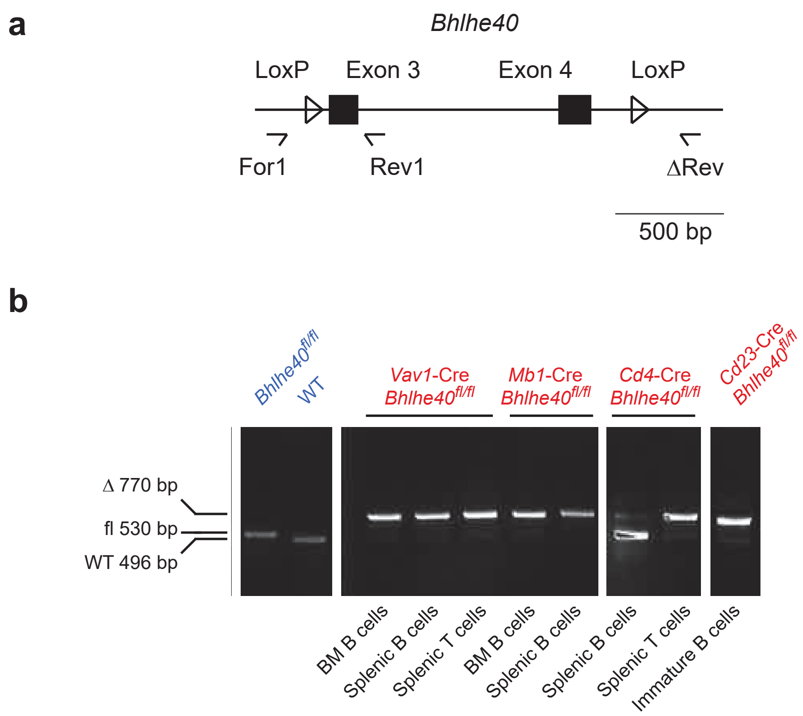
a
Bhlhe40
LoxP
Exon 3
Exon 4
LoxP
For1
Rev1
ΔRev
500 bp
b
Bhlhe40fl/fl
WT
Vav1-Cre Bhlhe40fl/fl
Mb1-Cre Bhlhe40fl/fl
Cd4-Cre Bhlhe40fl/fl
Cd23-Cre Bhlhe40fl/fl
Δ 770 bp
fl 530 bp
WT 496 bp
BM B cells
Splenic B cells
Splenic T cells
BM B cells
Splenic B cells
Splenic B cells
Splenic T cells
Immature B cells

Figure S6

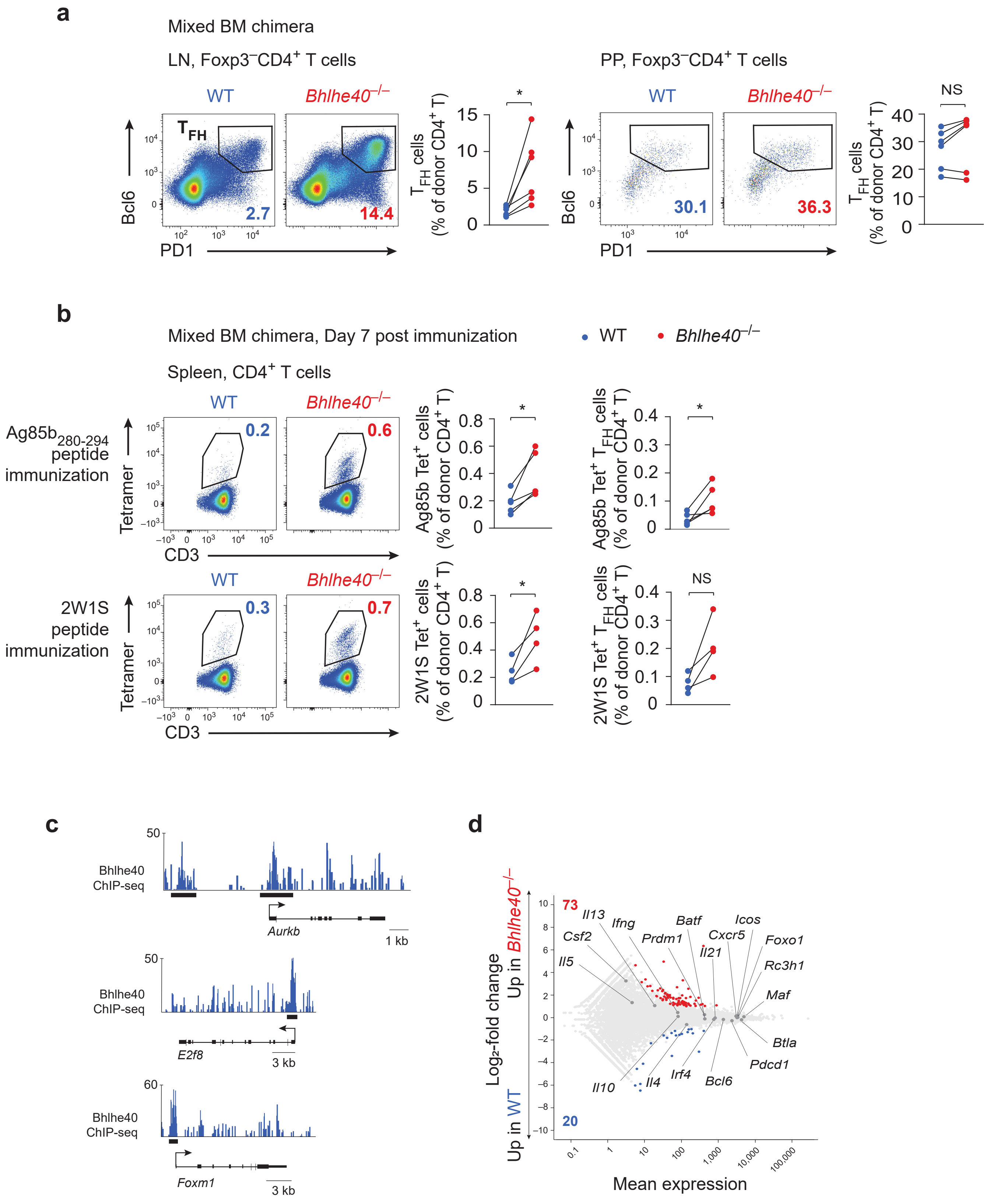
a
Mixed BM chimera
LN, Foxp3⁻CD4⁺ T cells
PP, Foxp3⁻CD4⁺ T cells
WT
Bhlhe40−/−
TFH
Bcl6
PD1
2.7
14.4
30.1
36.3
TFH cells (% of donor CD4+ T)
*
NS
b
Mixed BM chimera, Day 7 post immunization
WT
Bhlhe40−/−
Spleen, CD4+ T cells
Ag85b280-294 peptide immunization
2W1S peptide immunization
Tetramer
CD3
0.2
0.6
0.3
0.7
Ag85b Tet+ cells (% of donor CD4+ T)
Ag85b Tet+ TFH cells (% of donor CD4+ T)
2W1S Tet+ cells (% of donor CD4+ T)
2W1S Tet+ TFH cells (% of donor CD4+ T)
c
Bhlhe40 ChIP-seq
Aurkb
1 kb
E2f8
3 kb
Foxm1
3 kb
d
Log2-fold change
Up in Bhlhe40−/−
Up in WT
73
20
Il13
Ifng
Csf2
Il5
Prdm1
Batf
Icos
Cxcr5
Il21
Foxo1
Rc3h1
Maf
Btla
Pdcd1
Bcl6
Irf4
Il4
Il10
Mean expression

Figure S7

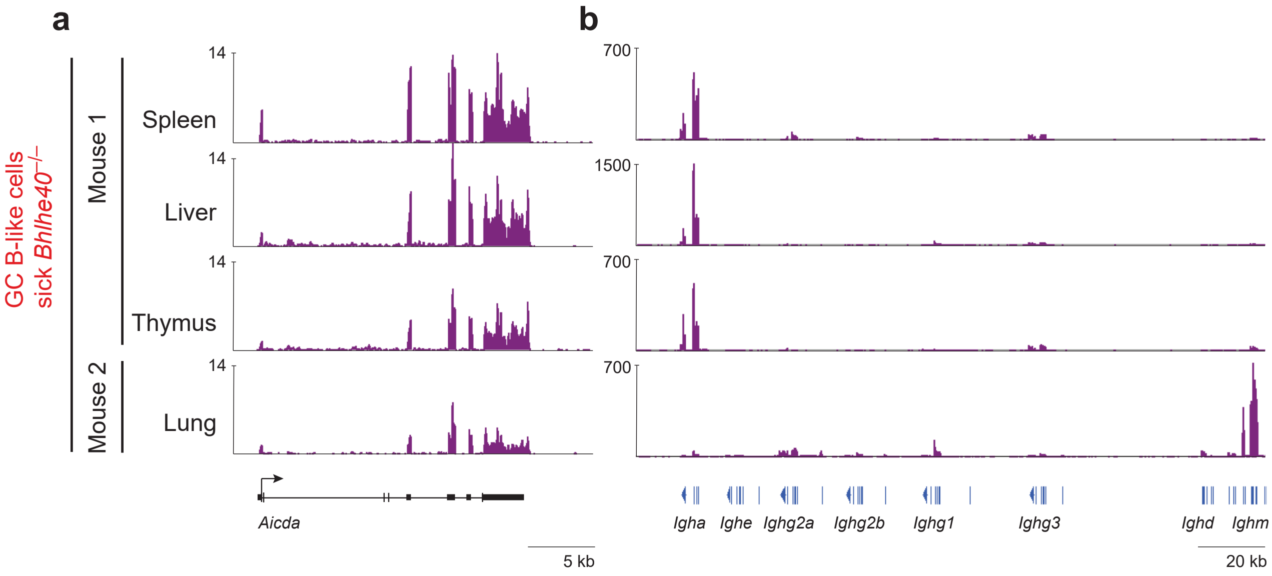
a
b
GC B-like cells
sick Bhlhe40−/−
Mouse 1
Mouse 2
Spleen
Liver
Thymus
Lung
14
700
1500
Aicda
5 kb
Igha
Ighe
Ighg2a
Ighg2b
Ighg1
Ighg3
Ighd
Ighm
20 kb

Figure S8

**Figure S1. Bhlhe40 restrains the GC reaction by a B cell-intrinsic mechanism.** (**a**,**b**) Spleen sections of unchallenged WT and *Bhlhe40*$^{-/-}$ mice as analyzed by confocal immunofluorescence microscopy. (**a**) The area of spontaneous GCs was determined by measuring the size of IgD-negative areas within B cell follicles. Pooled data with n = 2 mice for each genotype each are shown. (**b**) Spontaneous germinal centers (GCs), identified by PD1 (green) expression in IgD (red)-negative areas, are highlighted. One experiment is shown. (**c**,**d**,**f**) Mixed BM chimeras were generated by transferring a 1:1 (**c**,**f**) or 4:1 (**d**) mixture of congenically distinguishable WT (CD45.1$^{+}$) and *Bhlhe40*$^{-/-}$ (CD45.2$^{+}$) (**c**,**d**) or *Bhlhe40*$^{-/-}$*Bhlhe41*$^{-/-}$ (CD45.2$^{+}$) or *Bhlhe41*$^{-/-}$ (CD45.2$^{+}$) (**f**) BM progenitor cells into lethally irradiated *Rag2*$^{-/-}$ recipients. Mice were analyzed > 6 weeks after transfer. (**c**) Flow cytometric analysis and quantification of WT and *Bhlhe40*$^{-/-}$-derived cells among LSKs, CD19$^{+}$ B cells and CD4$^{+}$ T cells is shown. One experiment with n = 4 mice is shown. Representative of four independent experiments. (**d**) Representative images of spleen sections of 4:1 WT and *Bhlhe40*$^{-/-}$ mixed BM chimera. Spontaneous GCs, identified by GL7 (white) expression, and predominance of *Bhlhe40*$^{-/-}$-derived cells in the GCs is highlighted. WT-derived CD45.1$^{+}$ cells are shown in blue, *Bhlhe40*$^{-/-}$-derived CD45.2$^{+}$ cells in red. Note depletion of WT cells from the GC area. (**e**) Mixed BM chimeras were generated by transferring a 1:1 ratio of congenically distinguishable WT (CD45.1$^{+}$/CD45.2$^{+}$) and *Bhlhe40*$^{-/-}$ (CD45.1$^{+}$) BM progenitor cells into lethally irradiated WT (CD45.2$^{+}$) recipients. Quantification of flow cytometric analysis of GC B cells among WT and *Bhlhe40*$^{-/-}$ CD19$^{+}$ B cells is shown. One experiment with n = 5 mice, representative of two independent experiments, is shown. (**f**) Quantification of flow cytometric analysis of GC B cells in mixed BM chimeras among WT and *Bhlhe40*$^{-/-}$, *Bhlhe40*$^{-/-}$*Bhlhe41*$^{-/-}$ and *Bhlhe41*$^{-/-}$ CD19$^{+}$ B cells is shown. Pooled data from two independent experiments with n = 12 (WT and *Bhlhe40*$^{-/-}$), n = 8 (*Bhlhe40*$^{-/-}$*Bhlhe41*$^{-/-}$) and n = 4 (*Bhlhe41*$^{-/-}$) mice are shown. Representative of three independent experiments. Data were analyzed with Student's *t*-test (**a**, mean ± SD is shown), paired Student's *t*-test (**e**) or Wilcoxon matched-pairs signed-rank test (**f**). $^{**}P < 0.01$, $^{***}P < 0.001$. One dot represents one analyzed GC (**a**) or one mouse (**e**,**f**).

**Figure S2. Bhlhe40 expression in activated B cells.** (**a**) *Bhlhe40* expression in *in vitro* activated B cells is shown. Naive $CD43^{-}$ B cells from the spleens of WT mice were cultured up to 1 d in the presence of either anti-IgM or anti-CD40 antibody alone or in combination with IL-4. *Bhlhe40* expression (normalized to *Hprt*) was measured by RT-qPCR. Each data point represents the mean value with SEM of three technical replicates. Data are representative of two independent experiments. (**b**) *Bhlhe40* expression in follicular (FO) B and light zone (LZ) and dark zone (DZ) GC B cells. RNA-seq data from the Immgen database. (**c**) *Bhlhe40* expression in LZ and DZ GC B cells as assessed by RNA flow cytometry. Data are representative of two independent experiments. (**d**) Mixed BM chimeras were generated by transferring a 1:1 mixture of congenically distinguishable WT ($CD45.1^{+}$) and *Bhlhe40*$^{-/-}$ ($CD45.2^{+}$) BM progenitor cells into lethally irradiated *Rag2*$^{-/-}$ recipients. Mice were analyzed > 6 weeks after transfer. Flow cytometric analysis and quantification of the DZ/LZ ratio of WT and *Bhlhe40*$^{-/-}$ GC B cells are shown. One experiment with n = 5 mice is shown, representative of three independent experiments. Data were analyzed with paired Student's *t*-test. One dot represents one mouse.

**Figure S3. Effects of Bhlhe40 deficiency early during the GC response.** OVA-primed $CD45.2^{+}$ WT mice were injected with a 1:1 mixture of congenically distinguishable WT ($CD45.1^{+}$) and *Bhlhe40*$^{-/-}$ ($CD45.1^{+}$/$CD45.2^{+}$) (**a**) or WT ($CD45.1^{+}$/$CD45.2^{+}$) and *Bhlhe40*$^{-/-}$ ($CD45.1^{+}$) (**b**) B1-$8^{hi/+}$ splenocytes and were immunized with NP-OVA in alum next day. See schematic representation in Figure 2a. (**a**) Surface expression of CCR6 and GL7 on $CD45.1^{+}Ig\lambda^{+}CD19^{+}$ B cells on day 3 post immunization. Intracellular expression of Bcl6 in WT and *Bhlhe40*$^{-/-}$ $CCR6^{+}$ and GC B cells is shown. Representative staining for one of ten mice is shown; results representative of four independent experiments. (**b**) Surface expression of CCR6 and GL7 on $CD45.1^{+}Ig\lambda^{+}CD19^{+}$ B cells on day 3.5 post immunization. Representative flow cytometric analysis and quantification of WT and *Bhlhe40*$^{-/-}$-derived cells among $CCR6^{+}$ and GC B cells are shown. One experiment with n = 11 mice is shown, representative of at least three independent experiments.

**Figure S4. Bhlhe40 deficiency does not affect proliferation or apoptosis of activated B cells.** (**a**,**c**) Mixed BM chimeras were generated by transferring a 1:1 mixture of congenically distinguishable WT ($CD45.1^+$) and *Bhlhe40*$^{-/-}$ ($CD45.2^+$) BM progenitor cells into lethally irradiated *Rag2*$^{-/-}$ recipients. Flow cytometric analysis and quantification assessing the proliferation (**a**) and frequency of apoptotic cells (**c**) of light zone (LZ) and dark zone (DZ) (**a**) WT and *Bhlhe40*$^{-/-}$ GC B cells. (**a**) EdU was i.v. injected 2 hours prior to analysis. (**c**) Apoptotic cells were identified by intracellular staining for active caspase-3. (**a**,**c**) Pooled data from two independent experiments with n = 6 mice are shown. Data were analyzed with paired Student's *t*-test. One dot represents one mouse. (**b**) Naive $CD43^-$ B cells from the spleens of WT ($CD45.1^+$) and *Bhlhe40*$^{-/-}$ ($CD45.2^+$) mice were mixed in a 1:1 ratio and labeled with CTV. Cells were kept in culture for 3 d in the presence of decreasing concentrations of either anti-CD40 or anti-IgM antibody alone or in combination with IL-4. The ratio of WT and *Bhlhe40*$^{-/-}$-derived cells among live cells on day 3 of the culture and CTV dilution are depicted. One experiment with n = 3 mice is shown.

**Figure S5. Comparison of gene expression changes in WT and *Bhlhe40*$^{-/-}$ $CCR6^+$ and early GC B cells.** (**a**,**b**,**e**,**f**) OVA-primed $CD45.2^+$ WT mice were injected with a 1:1 mixture of congenically distinguishable WT ($CD45.1^+$) and *Bhlhe40*$^{-/-}$ ($CD45.1^+$/$CD45.2^+$) B1-8$^{hi/+}$ splenocytes and were immunized with NP-OVA in alum next day. RNA-seq analysis of double-sorted WT and *Bhlhe40*$^{-/-}$ $CCR6^+$ and early GC B cells was performed on day 3.5-4 post immunization. (**a**) Comparison of changes in gene expression induced by Bhlhe40 deficiency in $CCR6^+$ (top) and early GC B cells (bottom). $Log_2$-transformed *Bhlhe40*$^{-/-}$/WT fold changes are plotted. All significantly (> 2-fold, adjusted $P < 0.05$) changed genes are highlighted and the number of such up- and downregulated genes is indicated. (**b**) *Bhlhe40* expression in $CCR6^+$ and early GC B cells. (**c**) ChIP-seq analysis of Bhlhe40 binding in in vitro-activated B cells. Naive $CD43^-$ B cells from the spleen were activated in the presence of anti-CD40 for 4 h. (left) The consensus Bhlhe40-binding motif was identified by the *de novo* motif-discovery program. (right top) Example of Bhlhe40-binding at the *Bhlhe40* gene locus. (right bottom) Distribution of Bhlhe40 peaks at intergenic regions, the gene body or the promoter. (**d**) Bhlhe40 binding at the indicated gene loci (expression of the corresponding genes is shown in Figure 3c). Bars below the tracks represent called peaks. (**e**,**f**) UMAP plots as in Figure 3d highlighting the expression of CCR6$^{+}$-, GC B cell- and PB population-related genes (**e**) and GC B cell signature genes (**f**) in all cells in the data set.

**Figure S6. Deletion efficiencies of the loxP-flanked *Bhlhe40* allele.** (**a**) Schematic representation of the *Bhlhe40* floxed allele. (**b**) Deletion efficiency of the *Bhlhe40* allele in *Vav1*-Cre *Bhlhe40*$^{fl/fl}$, *Mb1*-Cre *Bhlhe40*$^{fl/fl}$, *Cd4*-Cre *Bhlhe40*$^{fl/fl}$ and *Cd23*-Cre *Bhlhe40*$^{fl/fl}$ mice as determined by PCR.

**Figure S7. Bhlhe40 restrains $T_{FH}$ cell numbers.** (**a**) Mixed BM chimeras were generated by transferring a 1:1 mixture of congenically distinguishable WT (CD45.1$^{+}$) and *Bhlhe40*$^{-/-}$ (CD45.2$^{+}$) BM progenitor cells into lethally irradiated *Rag2*$^{-/-}$ recipients. Mice were analyzed > 6 weeks after transfer. Expression of PD1 and Bcl6 by WT and *Bhlhe40*$^{-/-}$ Foxp3$^{-}$CD4$^{+}$ T cells and quantification of $T_{FH}$ cells among WT and *Bhlhe40*$^{-/-}$ Foxp3$^{-}$CD4$^{+}$ T cells from the lymph nodes (LN) and Peyer´s patches (PP) is shown. One experiment with n = 6 mice is shown, representative of two independent experiments. (**b**,**d**) Mixed BM chimeras were generated by transferring a 1:1 mixture of congenically distinguishable WT (CD45.1$^{+}$/CD45.2$^{+}$) and *Bhlhe40*$^{-/-}$ (CD45.1$^{+}$) BM progenitor cells into lethally irradiated WT (CD45.2$^{+}$) recipients. Mice were immunized with *M. tuberculosis* Ag85b$_{280\text{-}294}$ (**b**,**d**) or 2W1S (**b**) peptide emulsified in Complete Freund´s adjuvant (CFA) > 7 weeks after transfer. (**b**) Flow cytometric analysis and quantification of Ag85b- and 2W1S-tetramer-binding CD4$^{+}$ T cells and $T_{FH}$ cells among WT and *Bhlhe40*$^{-/-}$ CD4$^{+}$ T cells in the spleen on day 7 post immunization. For each condition, one experiment with n = 4 mice is shown, representative of one to two independent experiments. Data were analyzed with paired Student's *t*-test. *$P < 0.05$. One dot represents one mouse. (**c**) ChIP-seq analysis of Bhlhe40 binding at the indicated cell cycle-related gene loci in in vitro-activated CD4$^{+}$ T cells (data from a published dataset[42]). (**d**) RNA-seq analysis of Ag85b-tetramer-binding WT and *Bhlhe40*$^{-/-}$ CD4$^{+}$ T cells isolated from mixed BM chimeras 11 days post immunization. Comparison of changes in gene expression induced by Bhlhe40 deficiency in Ag85btetramer-binding $CD4^+$ T cells. $Log_2$-transformed *Bhlhe40*$^{-/-}$/WT fold changes are plotted and the number of significantly (> 2-fold, adjusted $P < 0.05$) up- and downregulated genes is indicated. $T_{FH}$ cell-related genes and selected cytokine-encoding genes are highlighted. Note that expression of none of these genes was significantly changed.

**Figure S8. RNA-seq analysis of GC B cell-like cells from sick *Bhlhe40*$^{-/-}$ mice.** mRNA expression profile at the *Aicda* (**a**) and *Igh* (**b**) locus in GC B cell-like cells of two sick *Bhlhe40*$^{-/-}$ mice as analyzed in Figure 6h.
